# Separate attentional processes in the two visual systems of jumping spiders

**DOI:** 10.1101/2023.04.13.536553

**Authors:** Federico Ferrante, Maria Loconsole, Davide Giacomazzi, Massimo De Agrò

## Abstract

By selectively focusing on a specific portion of the environment, animals can solve the problem of information overload, toning down irrelevant inputs and concentrate only on the relevant ones. This may be of particular relevance for animals such as the jumping spider, which possess a wide visual field of almost 360° and thus could benefit from a low-cost system for sharpening attention. Jumping spiders have a modular visual system composed of four pairs of eyes, of which only the two frontal eyes (i.e., AMEs) are motile, whereas the other secondary pairs remain immobile. We hypothesized that jumping spiders can exploit both primary and secondary eyes for stimulus detection and attentional shift, with the two systems working synergistically. In Experiment 1 we investigated AMEs’ attentional responses following a spatial cue presented to the secondary eyes. In Experiment 2, we tested for enhanced attention in the secondary eyes’ visual field congruent with the direction of the AMEs’ focus. In both experiments, we observed that animals were faster and more accurate in detecting a target when it appeared in a direction opposite to that of the initial cue. In contrast with our initial hypothesis, these results would suggest that attention is segregated across eyes, while each system works to compensate the other by attending to different spatial locations, rather than sharing a common attentional focus.

## 2 Introduction

Animals are continuously bombarded with sensory input, to the point that it would be impossible with their limited cognitive resources to compute and respond to each of them. With respect to visual attention, evolution favoured the emergence of a set of mechanisms that allow animals to selectively focus on specific stimuli or portions of the environment that might be relevant to them, while ignoring those that are irrelevant (Evans et al., 2011). Analogous attentional mechanisms appear to be widespread among different species and even clades, including mammals (Bowman et al., 1993; Marote and Xavier, 2011; Rizzolatti, 1983; Wagner et al., 2014), birds (Johnen et al., 2001; Lazareva et al., 2012; Shimp and Friedrich, 1993; Sridharan et al., 2014), amphibians (Greenfield and Rand, 2000; Tárano, 2015), fishes (Gabay et al., 2013; Parker et al., 2012) and even invertebrates (Eckstein et al., 2013; Humphrey et al., 2018; Morawetz and Spaethe, 2012; Wiederman and O’Carroll, 2013; Winsor et al., 2021).

Although individuals are generally in control of which stimuli to focus on, there are instances in which this process is forcibly interrupted and attention is automatically redirected toward a different location (Kouider and Dehaene, 2007). This happens, for example, in the presence of an unexpected stimulus with a relatively high intensity, such as a brief flash of light. In such cases, the subject automatically redirects attention to the source of the new stimulus (Posner, 2016). This allows individuals to quickly react to unexpected environmental changes. The spatial cuing task (Posner et al., 1980), or Posner task as per its inventor name, has long been used in the investigation of automatic redirection of attention in humans and other vertebrates (Bowman et al., 1993; Bushnell et al., 1981; Golla et al., 2004; Marote and Xavier, 2011; Wagner et al., 2014). The classical paradigm consists of a visual detection task in which subjects are required to locate a target stimulus. Before the target, a task-irrelevant cue is presented, either in the same location where the target will appear or in a different one (i.e., valid vs. invalid trials). If the task-irrelevant cue causes an automatic and involuntary shift of the attentional focus toward the location where it appeared, we expect subjects to be faster and more accurate in locating the target in valid trials compared to invalid trials. By varying the characteristics of either the cue or the target, researchers described differences in automatic attentional shift processes (Posner et al., 1980). In particular, Posner’s paradigm revealed that shifting attention does not necessarily require moving the visual focus (i.e., the eyes). This latter case would be an example of ‘covert’ attention, where the individual is more activated and prepared to respond to stimuli in a certain spatial location, despite the fact that their eyes are oriented toward a different portion of the space. (Theeuwes et al., 1998). On the other hand, an ‘overt’ attentional shift (Yantis and Jonides, 1984) refers to when individuals actively move their eyes toward the location of interest. These two are distinct mechanisms (Hunt and Kingstone, 2003a; Hunt and Kingstone, 2003b).

During the last decades, jumping spiders established themselves as a deeply interesting model for the study of vision-based behaviour, and have been found capable of producing complex behaviours (Cross et al., 2020; De Agrò, 2020; De Agrò et al., 2017; De Agrò et al., 2021; Dolev and Nelson, 2014; Dolev and Nelson, 2016; Harland and Jackson, 2000; Rößler et al., 2022a; Rößler et al., 2022b). On the one hand, it is possible to hypothesise that jumping spiders could rely on attentional visual mechanisms analogous to those of vertebrates that similarly rely on vision as the primary system. On the other hand, jumping spiders could resort to a different attentional mechanism, due to their peculiar brain organization. These arthropods in fact possess a modular visual system (Foelix, 2011; Harland and Jackson, 2000; Land, 1985; Menda et al., 2014; Morehouse, 2020; Zurek and Nelson, 2012; Zurek et al., 2010), which charges upon 4 different eye pairs. Large anteromedial eyes (AME, “primary eyes”) are characterised by a narrow visual field (≈5° for each eye), while having a high visual acuity and being capable of colour vision (Blest et al., 1981; Foelix, 2011; Jackson and Cross, 2011; Land, 1969a; Land, 1969b; Morehouse, 2020; Zurek et al., 2015). The long eye tubes of these primary eyes can be moved by the spider thanks to a set of muscles (Land, 1969b). These characteristics suggest that the AME are specialised in the detection of fine details and figure recognition (Harland et al., 2012). The other 3 pairs of eyes, anterolateral, posteromedial, and posterolateral eyes (ALE, PME, PLE, ‘secondary eyes’), are unmovable, monochrome, and are characterised by a lower visual acuity but a much wider visual field, covering a combined ≈350° around the spider. These eyes are specialised in motion detection and recognition (Bruce et al., 2021; De Agrò et al., 2021; Duelli, 1978; Fenk and Schmid, 2010; Jackson and Cross, 2011; Jakob et al., 2018; Spano et al., 2012; Zurek and Nelson, 2012; Zurek et al., 2010). The different eyes project into separate networks (Steinhoff et al., 2020), and are seemingly independent from each other, apart from late multimodal integration (Strausfeld and Barth, 1993; Strausfeld et al., 1993). This division of tasks is also reflected in the spider’s behaviour: if a target falls in the secondary eyes’ visual field, the spider rotates its whole body until it points the AME directly towards the detected object (Harland and Jackson, 2000; Zurek et al., 2010). These whole-body pivots are termed ‘saccades’, due to the functional parallelism with the homonymous behaviour in humans. Once the target is locked on by the AME, their long eye tubes start moving, scanning the object, and following it in case of dislocation (Jakob et al., 2018; Land, 1969a).

During AME scanning, the secondary eyes remain vigilant and can still detect novel stimuli entering their visual field. Bruce et al. (2021) showed that if a stimulus is presented in the peripheral visual field of the spider while its attention is overtly focused on a target (i.e., the spider is pointing the primary eyes), the animal would disengage from the target to redirect the primary eyes toward the new stimulus. Such a behaviour somehow mirrors the human Posner effect in that attention is forcibly shifted toward a second portion of the space because of an unexpected salient stimulus appearing out of the overt focus. It remains unclear whether the secondary eyes themselves possess a separate, covert, attentional mechanism or if they only rely on the overt AME attentional shift to inform computational precedence.

Following this parallelism with Posner’s attention task, we investigated whether a brief spatial cue in the peripheral visual field (that is, presented to the secondary eyes) of the spider could pre-alert and increase covert attention toward that portion of the space, resulting in a facilitation in locating a target congruent with the position signalled by the cues rather than a target appearing in the opposite hemispace. In a first experiment, we presented jumping spiders with a spatial cue (i.e., a brief flash of light) to the right or left side on a screen, followed by a target stimulus (i.e., a white dot moving vertically) in the same or opposite position. If the presentation of the cue could enhance the attentional system via the secondary eyes, and these could synergistically pre-alert the primary eyes, we could hypothesise a reduction in reaction speed or an increase in body saccade frequency toward the target in valid (i.e., target appearing in the same location as the cue) versus invalid (i.e., target appearing in the opposite location as the cue) trials. In a second experiment, we investigated whether orienting the primary eyes overt attention toward a certain portion of the space could favour the secondary eyes detection of stimuli appearing in the nearby areas of the AME visual field, further supporting our initial hypothesis of a shared attentional system between the primary and secondary eyes.

## 3 Methods

### 3.1 Subjects

We employed a total of 136 *Menemerus semilimbatus*. 65 spiders participated in Experiment 1, and the remaining 71 in Experiment 2. All animals were collected in nature, in the park outside the Esapolis’ living insects museum, Padua, Italy, between March and September 2021. Once caught, we applied to the cephalothorax a cylindrical, 1×1mm neodymium magnet, using UV activated resin. This magnet was needed to fix the animal to the apparatus (see the next section). The spiders were then housed individually in plastic boxes and tested for the first time the next day. At the end of the experiment for each animal, the magnet was removed and the subject was released back into nature.

### 3.2 Overall procedure

Both experiments were performed using a spherical treadmill, as fully described in (De Agrò et al., 2021). The spider was attached to the end effector of a 6-axis micro-manipulator. The magnet glued to the spider’s cephalothorax would connect to an opposite-polarity magnet on the end effector. Just below, a polystyrene sphere was present. This ball was maintained suspended by constant streams of compressed air, to make its motion frictionless. By adjusting the micromanipulator, the spider was approached to the sphere, until it got in contact with its legs. In this configuration, the spider orientation would remain fully fixated, while it could still impress its intended movements to the sphere. By recording and extracting the rotation of the sphere, using the software FicTrac (Moore et al., 2014), we could infer animal behaviour. The full apparatus was put at 250mm of a computer screen, directly in front from the spider’s perspective. The monitor had a 1080p resolution, a refresh rate of 60hz, and a pixel pitch of 0.248mm. The stimuli were presented on the monitor at 30fps, at 30° left or right from the center of the visual field. The recording and the stimuli presentation were programmed in Python 3.8 (Van Rossum and Drake, 2009), using the packages psychopy (Peirce et al., 2019), opencv (Bradski, 2000), numpy (Oliphant, 2006; van der Walt et al., 2011). As stated in the introduction, jumping spiders perform peculiar full-body saccades towards a target perceived in the secondary eyes visual field, to focus it with the primary eyes. By observing the Z-axis rotations of the sphere, we were able to extract intended spiders’ rotations (i.e., body saccades) towards each stimulus.

### 3.3 Experiment 1 - Attentive priming with peripheral cues

In our first experiment, we inquired about the effect of a peripheral cue presented in the secondary eyes’ visual field on the speed and probability of detection of target by the same eyes. To start, the background of the monitor was set to gray, apart from two “windows”, at ±30° from the centre, respectively. These windows had a white background and were 20° wide and 20° tall (Figure 1).

**Fig. 1:**
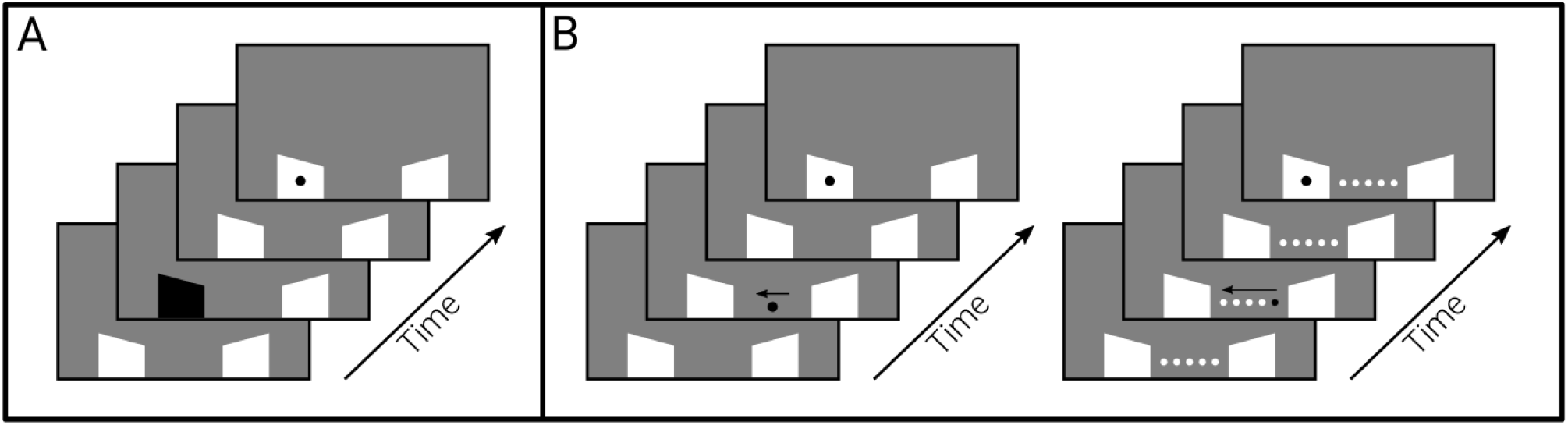
Experimental procedure. **A)** First experiment setup. The peripheral cue consisted of a brief flash on one of the two windows, alternating between black and white for 0.2 seconds at a speed of 30hz. After the cue disappearance, a random time interval between 0.2 and 5 seconds would elapse before the target appeared in the same or opposite position. **B)** Second experiment setup. In the “moving dot” condition, the cue stimulus, a wide circle, appeared at the center of the screen moving towards one of the two windows. In the “five dots” condition, five dots at the center of the visual field simulated the movement towards one of the windows by alternating their luminance from black to white sequentially, before the appearance of the target.

**Fig. 2:**
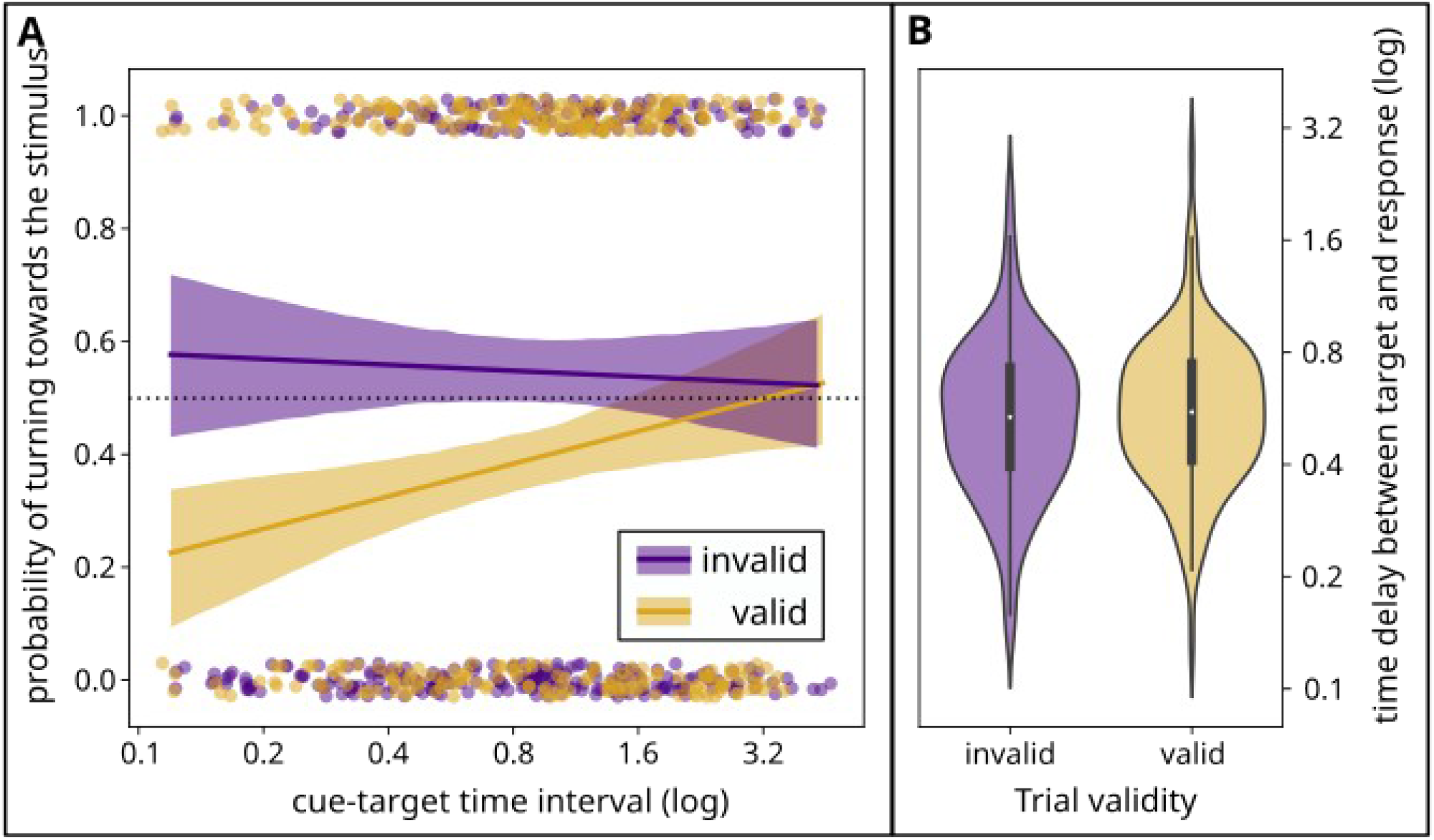
First experiment results. **A)** Probability of turning towards the target stimulus in valid vs invalid trials. The X-axis represents the logarithmic interval (in seconds) between the cue disappearance and the target stimulus appearance. The Y-axis represents the response probability, as the average probability that subjects respond to the appearance of the target stimulus (in percentage). **B)** Response time to the target stimulus, as the time between the target appearance and the beginning of a response towards the target. On the X-axis the valid vs invalid conditions, the Y-axis represents the interval (in seconds) between target and response.

**Fig. 3:**
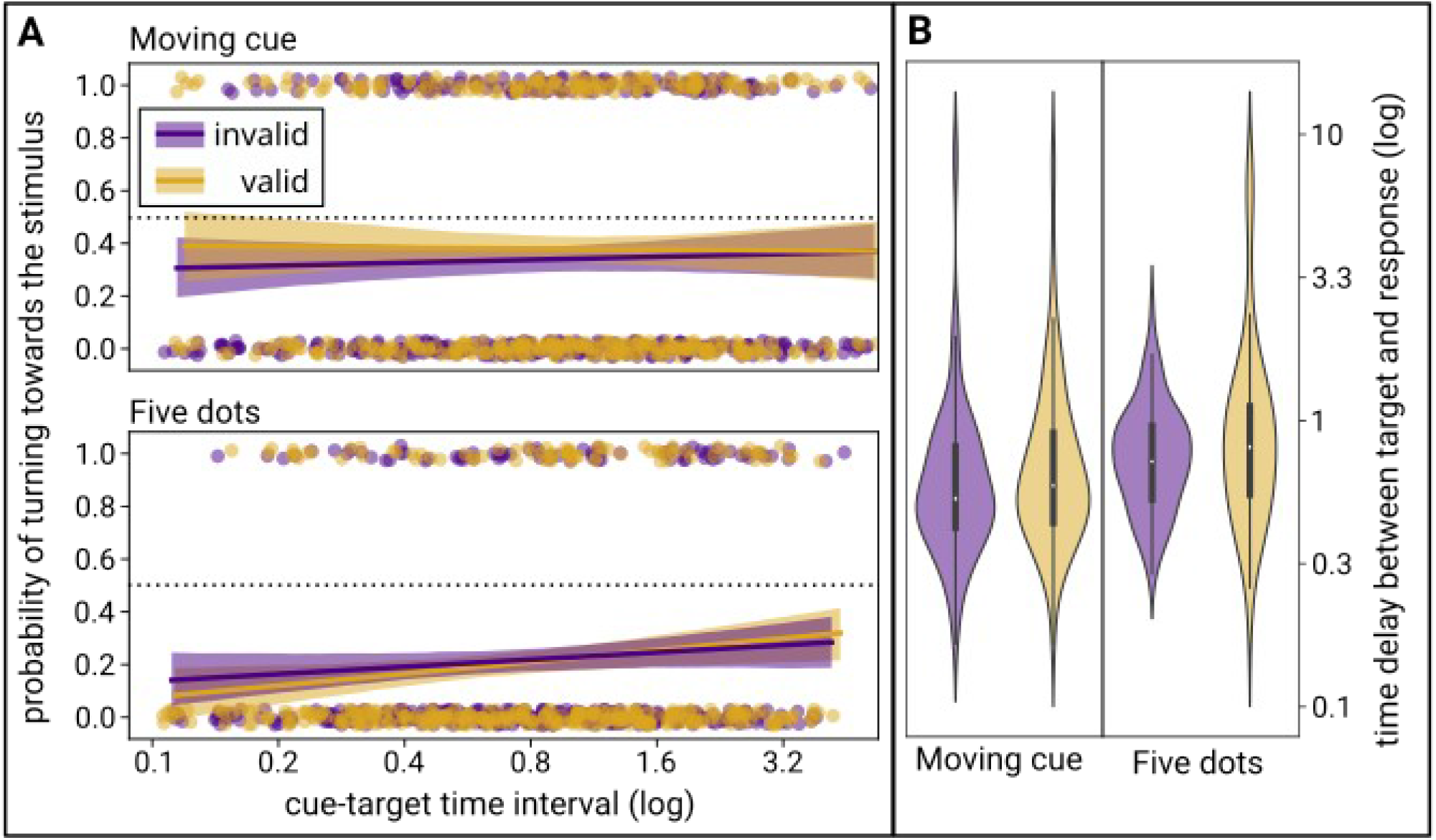
Second experiment results. **A)** Probability of turning towards the target stimulus in “moving cue” and “Five dots” conditions (valid vs invalid trials). The X-axis represents the logarithmic interval (in seconds) between the cue disappearance and the target stimulus appearance. The Y-axis represents the response probability, as the the average probability that subjects respond to the appearance of the target stimulus (in percentage). **B)** Response time to the target stimulus, as the time between the target appearance and the beginning of a response towards the target. On the X-axis the valid vs invalid trials for both the conditions. The Y-axis represents the interval (in seconds) between target and response.

After 3 minutes of habituation, in which the spiders observed this static background while harnessed to the treadmill apparatus, the first stimulus appeared. One of the two windows would flash”, alternating between black and white for 0.15 seconds at a speed of 30hz. This flash would act as a spatial cue, aiming to increase the spider’s covert attentional monitoring toward that portion of the space. After the cue disappearance, a random time interval between 0.2 and 5 seconds would elapse. Randomisation was performed logarithmically: larger intervals were sparser than shorter ones. This was done on the assumption that the interval effect is logarithmic itself, as many psychophysical effects (Portugal and Svaiter, 2011). After this interval, the target stimulus was presented: this consisted of a 4° wide circle, moving up and down in alternating frames for a span of 2°, for a total of 0.66 seconds (Figure 1A). This target could appear at the centre of one of the two lateral windows, either the one previously pre-alerted by the cue (i.e., valid trials) or the opposite one (i.e., invalid trials). After the target disappeared, a 25 seconds pause was presented, followed by the next cue-interval-target set. In a full session, the spiders were presented with a total of 12 trials.

We also designed two control conditions. In the first condition, the spiders were presented directly with the target stimulus, without being preceded by the flashing light cue. This was used to collect a baseline measure of response rate and speed. In the second condition, the spiders were presented with the cue only, without being followed by a target. This was used to check whether the cue alone was sufficient to elicit saccades. It is possible in fact that in the main condition, the spiders would perform a body saccade towards the cue but slow enough to be mistakenly recorded as a fast reaction to the subsequently presented target. By testing the reaction to cue alone, we could control for this possibility. The trials of these two conditions also started with the 3 minutes of habituation and presented a total of 12 trials. Each spider was subjected to all 3 conditions, in 3 different sessions, on the same day.

### 3.4 Experiment 2 - Attentive priming with central cues

In our second experiment, we investigated whether directing the AME attention toward a spatial location could favour stimulus detection in the secondary eyes’ visual field in that same portion of the space. To this aim, we presented a central spatial cue in the AMEs visual field. This was designed to trigger a visual shift in the AMEs, either toward the left or right. We designed two different versions of such cues for two separate experimental conditions. Both conditions maintained the same overall structure as the first experiment (background color, window position, initial habituation section).

In the first condition, the cue, named ‘moving dot’, consisted of a 4° wide circle, appearing in the centre of the screen, and moving toward one of the two windows, for a span of 15°, at a speed of 15°/sec, to finally disappear again (Figure 1B).

In the second condition, the employed cue was named ‘five dots’. It consisted of 5 dots at the centre of the visual field. The dots were equally spaced, 7.5° from each other, ranging from -15° to +15°. The dots had a diameter of 4° and were white in colour. These 5 dots remained visible on the screen for the whole duration of the experiment. The cue consisted of the change in colour of the left or right circle, from white to black. After 0.15 seconds, the dot switched back to white and the adjacent dot turned black. This proceeded until the black dot reached the opposite side, leaving only white dots. Notably, AMEs visual field covers only 10° (5° for each side to the centre); each dot appeared at 7.5° degrees distance from the previous one, forcing the spider to use the ALEs for detection and then re-anchor the AMEs focus onto it (Figure 1B). This created a substantial difference with respect to the moving dot condition, where the spider could follow the target with the AMEs alone, as it moved for less than 1° per frame, always remining in focus. Moreover, differently from the first condition, the first dot changing colour was not centred but already oriented in space (i.e., either the left or the right one). As such, the probability of the AME starting with the dot in the visual field decreased even more. Overall, this condition was intended to maintain the same indication towards one of the two sides of the previous one, but without stimulating the AMEs, forcing the spiders to rely on the ALEs fields.

In both conditions, following a random time interval (see Experiment 1), a target appeared either in the window on the same side of the cue disappearance (i.e., valid trial) or the other one (i.e., invalid trial). We expected the cue to be tracked by the spider’s AMEs, causing a shift of the animal’s overt attention to a specific direction. Importantly, it was not possible for the spider to reach the two lateral windows with the AMEs, as they were positioned outside of their visual fields. As such, any stimulus appearing in the two lateral windows had to be detected relying on the secondary eyes.

### 3.5 Scoring

The rotations of the spherical treadmill, collected with the software FicTrac (Moore et al., 2014), were further analysed through an algorithm developed in Python 3.8 (Van Rossum and Drake, 2009), available as a supplement (ESM02).

We analysed z-axis rotations in the 5 seconds after the appearance of each stimulus (i.e., the cure or the target). Cumulative changes in absolute orientation of at least 20° (the angle was chosen to be consistent with the stimulus position at 30°, but still accounting for errors) in a direction consistent with the position of the stimulus (e.g., if the stimulus was to be located on the left, we would expect clockwise rotations of the sphere) were considered as body-saccades toward that stimulus.

We also scored the latency of movement by calculating the time that elapses between the appearance of the stimulus and the beginning of the rotation. For what concerns Experiment 1, in the ‘only cue’ condition, the latency was calculated from the presentation of the cue, whereas in the two other conditions it was calculated from the presentation of the target. In these latter cases, we also checked for rotations occurring from the presentation of the cue up to 5 seconds after the target appearance. Regarding Experiment 2, we calculated latency only following the presentation of the target.

### 3.6 Statistical analysis

All analyses were performed in R4.1.2 (R Core Team, 2021), using the packages readODS (Schutten et al., 2020), glmmTMB (Bolker et al., 2009; Brooks et al., 2017), car (Fox and Weisberg, 2019), DHARMa (Hartig, 2018), emmeans (Lenth, 2018), ggplot2 (Wickham, 2009) and reticulate (Ushey et al., 2021). Raw data is available in ESM01.

For the first experiment, we initially observed the differences between the three conditions, both in terms of response probability and response delay. For the saccade probability, we employed a generalised linear mixed model (GLMM) with a Binomial error structure. We also included session and trial number, as we expected to observe a decrease in saccades with the passing of time, as previously suggested (Melrose et al., 2018). We included the identity of the subjects as a random intercept, and trial number as a random slope. After evaluating the goodness of fit for the model, we observed the effect of the predictors using an analysis of deviance, and subsequently performed a Bonferroni-corrected post hoc analysis on factors that had a significant effect. When analysing the response delay, we followed the same procedure, but used a Gaussian error structure. Notably, the analysis was performed on the delays log, rather than on the raw value. This decision followed the same logic we used when deciding the time intervals between cues and targets: in the realm of psychophysics, the magnitude of change increases logarithmically, rather than linearly (Portugal and Svaiter, 2011 see ESM2 for the distributions).

After we had observed the difference between conditions, we concentrated on the effects in the main experimental condition, where both the cue and the target were presented. We again analysed both the probability of response and the response delay, using the GLMM binomial and Gaussian error structures, respectively. As predictors, we included trial validity (cue in the same or opposite spatial location of the target) and time interval (between the appearance of cue and the one of target), in addition to trial and stimulus number as before. We included the identity of the subjects as a random intercept, and stimulus number as a random slope. The analysis then followed the same procedure described above.

The analysis for the second experiment followed the exact same structure, observing as dependent variables both the probability of saccades and the response delay. For both models, we included as predictor the experimental condition (‘moving dot’ or ‘five dots’), the validity of the trial, the time interval and the stimulus number (the trial number was not included because it was shown to have no effect for this experiment, see ESM02).

## 4 Results

### 4.1 Experiment 1 - Attentive priming with exogenous cues

As expected, we found an overall effect of both the trial number (GLMM analysis of Deviance, χ2=13.939, DF=1, p=0.0002) and the session number (χ2=32.947, DF= 2, p<0.0001) on the probability of producing a saccade toward any stimulus. Specifically, the response probability remains the same in sessions 1 and 2 (corrected post hoc Bonferroni, est = 0.0139, SE = 0.124, t = 0.121, p = 1), while for session 3 it is lower than the other 2 (1vs3, est.=0.707, SE=0.138, t=5.131, p<0.0001; 2vs3, est.=0.0.693, SE=0.137, t=5.07, p<0.0001). Across trials in each session, response probability significantly decreases (trend=-0.056, SE=0.0153, t=-3.633, p=0.0003).

Regarding the differences between the three conditions, the response probability across all trials was the same for the ‘only target’ condition and the ‘both cue and target’ condition (post-hoc Bonferroni corrected, odds ratio=1.0656, SE=0.274, t=0.247, p=1). On the contrary, both conditions presented a higher response probability relative to the ‘only cue’ condition (both cue and target VS only cue, odds ratio=31.585, SE=10.7, t=10.191, p<0.0001; only cue VS only target, odds ratio=0.0337, SE=0.0116, t=-9.823, p<0.0001). Similarly, the conditions of ‘both cue and target’ and ‘only target’ were characterized by a faster response delay over the ‘only cue’ condition (‘both cue and target’ VS ‘only cue’, ratio=0.151, SE=0.0164, t=- 17.347, p<0.0001; ‘only cue’ VS ‘only target’, ratio=5.075, SE=0.5933, t=13.897, p<0.0001). The condition ‘both cue and target’ also had faster response rate than the ‘only target’ condition, although the difference was much smaller (ratio=0.765, SE=0.0709, t=-2.885, p=0.0113). These results suggest that cue alone does not trigger a saccade, for which the target is required instead.

Regarding the response delay in the main condition, we found no effect of the time between the cue and the target appearance (GLMM analysis of Deviance, χ2=0.3866, DF=1, p=0.534). Instead, there was an effect of trial validity (χ2=7.9906, DF=1, p=0.0047). Specifically, spiders showed faster responses (0.505 seconds vs. 0.584 seconds) during invalid trials over valid ones (post-hoc Bonferroni corrected, ratio=0.866, SE=0.0432, t=-2.894, p=0.0041).

The response probability was strongly influenced by both the validity of the trial (GLMM analysis of Deviance, χ2=17.8217, DF=1, p<0.0001) and by the interaction between trial validity and the time between the cue and target appearance (χ2=5.2378, DF=1, p=0.0221). Specifically, the response probability was higher for invalid trials than for valid ones (post hoc Bonferroni corrected, odds ratio=2.58, SE=0.628, t=3.889, p=0.0001). Furthermore, while the response probability remained constant over time intervals for invalid trials (trend=-0.108, SE=0.157, t=-0.69, p=0.4905), it increased significantly for valid ones (trend=0.471, SE=0.165, t=2.853, p=0.0045), starting from a much lower response for short intervals and reaching an equal probability of invalid trials for longer ones.

### 4.2 Experiment 2 - Attentive priming with endogenous-like cues

For the second experiment, we found no differences between the two sessions in terms of response probability (GLMM analysis of Deviance, χ2=2.61, DF=1, p=0.106). This is consistent with the result of experiment one, as the difference observed there was only evident in session 3. We also found an overall effect of the trial number independently of the condition (χ2=76.497, DF=1, p<0.0001), as previously observed.

Regarding the response delay in both conditions, we found an overall effect of the validity of the trial (GLMM analysis of Deviance, χ2=4.756, DF=1, p=0.0292). Instead, there was no effect of the time interval between the cues and targets (χ2=0.8371, DF=1, p=0.3602), nor did we observe any differences between the two conditions (χ2=0.651, DF=1, p=0.4197). Specifically, spiders appear to be marginally faster for invalid trials than for valid ones (post hoc, ratio = 0.87, SE = 0.0625 t = -1.941, p = 0.053), although post hoc does not report significance as the analysis of deviance does due to the Bonferroni correction, suggesting that the effect is small.

Regarding the probability of response, we found a difference between the two conditions (GLMM analysis of Deviance, χ2=15.618, DF=1, p<0.0001), but no effect of trial validity (χ2=0.6137, DF=1, p=0.433) nor of the time interval between cue and target (χ2=3.5913, DF=1, p=0.058). Specifically, spiders have a much higher probability of responding to targets in the ‘moving dot’ condition over the ‘five dots’ one (post hoc, odds ratio=0.254, SE=0.0784, t=-4.482, p<0.0001).

## 5 Discussion

In this paper, we enquired the presence of a covert visual attentional mechanism in jumping spiders, similar to what reported in humans, and whether this relies on a synergistic activation of both primary and secondary eye systems.

In the first experiment, we investigated whether a spatial cue presented to the secondary eyes could affect the detection of a target stimulus also presented to the same eyes. According to the literature on vertebrates (Posner, 2016; Posner et al., 1980), we hypothesised that a spatial cue would support the performance when appearing in the same position of the target (valid trials) and hinder it when appearing in the opposite hemispace (invalid trials), both in terms of probability of detection and response times. We observed, contrary to our expectations, that subjects performed better in invalid trials, where the probability of detecting the target remained stable across time intervals and spiders were overall faster in responding compared to valid trials. In these latter, the probability of response was affected by the time interval, with low scores for short intervals (around 0.2 seconds) and an increasing improvement until the performance of the invalid trials was matched in longer delays (around 5 seconds).

The initially unexpected drop in performance for valid trials could be explained with the peculiar split role of spider AMEs and ALEs, and the possibly split nature of attention between these two sets of eyes. In natural behaviour, following a first detection by the ALEs, there should be an overt shifting (i.e., the spider performs a saccade to orient toward the stimulus) of the AMEs. These anchor to the stimulus detected by the ALEs and start a detailed scan. It is possible to hypothesise that, to maximise the efficiency of the allocation of attentional resources, after stimulus detection the ALEs will dis-anchor from the previously detected stimulus and redirect their (covert) attention to a different portion of the space. The ALEs would then be inhibited from re-anchoring on the previously attended hemispace, in order to avoid an overlap with the AMEs currently attended area, maximizing the efficiency and ability to detect novel stimuli. In the case of our experiment, this resulted in a worsening of detection in valid trials (both in terms of probability of detection and reaction times, as we observed), while maintaining an optimal performance in invalid ones, as ALEs would still be responsive to portions of the space other than the cue’s. Note that when the spiders do respond to the targets, they are significantly faster with respect to when no cue was presented at all, and consequentially no covert attentional shift of ALEs occurred. This shows that shutting off a portion of the visual field is indeed advantageous for performance. This ALEs attentional blindness with respect to the previously inspected spatial location has a limited duration, as the performance is restored after longer delays between the cue and the target, in the order of 5 seconds.

In the ‘moving dot’ condition of the second experiment, we addressed whether a translating visual cue presented to the AMEs could improve detection of a spatial target in the visual field of the secondary eyes. The cue appeared in the centre of the screen, where it could be detected by the AMEs. We know from previous literature that smooth tracking of a target with AMEs is directed by the ALEs (Zurek and Nelson, 2012). As such, while the cue moved across the screen, we can assume that both AMEs’ and ALEs’ attention shifted with it. Similarly to Exp. 1, spiders responded faster to invalid trials rather than to valid ones. This is again contrary to our initial hypothesis, yet it well fits our interpretation of the results in terms of the ALEs being hindered in maintaining visual attention in a previously inspected spatial location. In terms of reaction times, spiders were overall faster in invalid *versus* valid trials, hinting at an enhanced attention in the previously unattended location (or, conversely, a reduced attention in the already inspected one). However, while this effect was found to be significant in the ANOVA, it was lost in the post hoc analysis. This is probably due to the low response rate in Exp. 2, for which, despite the number of subjects being equal, we had fewer observations to be included in the analysis.

Differently from Exp. 1, when considering the probability of response, we did not find a difference between valid and invalid trials. This could be explained as a result of methodological differences between the two experiments. In fact, in Experiment 1 the target would appear at exactly the same location as the cue, hence being affected by the hypothesised attentional blindness of the ALEs; however, in Experiment 2, valid trials presented the target stimulus in the same direction as the cue (left or right) but not in the exact same spatial position. As such, the target stimulus could be still considered as novel rather than completely overlapping with the cue. As such, it could have been deemed relevant and attended, even if with a slower reaction time due to the attentional shift.

Interestingly, we found a difference between the ‘moving dot’ and the ‘five dots’ conditions, the latter scoring significantly lower response rate. We interpreted this as due to a different involvement of the primary and secondary eye systems. In the ‘moving dot’ condition, as stated before, the AMEs could smoothly track the cue to the left or right hemispace. Therefore, we can assume that the ALEs’ attentional focus moved together with the AME’s toward the cued direction, in a process that followed these steps: i) first detection of the cue by the ALEs; ii) subsequent involvement of the AMEs for sustained attention; iii) attentional tracking of the moving cue from both the AMEs and the ALEs; iv) second detection of the target by the ALEs. However, in the ‘five dots’ condition the cue could not be smoothly followed, as it did not proceed along a continuous movement, but rather haltingly appeared and disappeared, progressively shifting toward a certain spatial location. Compared to the ‘moving dot’ condition, this would require repeated effort to detect the dot reappearing in the novel location and re-anchoring the attention to a newly appeared object for a total of six times for each trial (five times for the five dots acting as a cue, plus one additional time for the target). This could constitute a highly demanding task for the spiders, resulting in the observed drop in response rate. This effect would also be enhanced by the hypothesised re-anchoring refractoriness to previously attended hemispaces. Similarly, we observed a reduction in the response rate between trials within the same session in both our experiments. This is also in line with previous literature showing that jumping spiders’ detection rate drops following repeated stimulus presentation (De Agrò et al., 2021; Humphrey et al., 2018; Humphrey et al., 2019; Melrose et al., 2018).

Overall, our study hints at the presence of two independent systems for visual attention, for which the jumping spider’s primary and secondary eyes work independently, but synergistically, in attentional monitoring and stimulus detection. When the ALEs detect a stimulus in space, the AMEs start a scanning process of it. This would free attentional resources for the ALEs with respect to that portion of space, allowing for monitoring different locations and not overlapping with the AMEs attentional focus. In the case of our study, it was not possible for spiders to coordinate the two systems’ attentional focus, as the animals were constrained on a fixed position on the sphere, leading to the discussed poor performance in valid trials, as the AMEs could not overtly direct attention to the cue location while the ALEs were covertly scanning different portions of the space. However, at the same time, we could observe an advantage in invalid trials (Exp. 1) for which spiders were faster in responding to the target when it was followed by the cue compared to when the cue was not presented. This is consistent with the idea that the spider narrows the attentional focus to spatial locations that were not previously inspected. Indeed, even though the split attention mostly hindered the spider performance in our experiment, it would probably be highly advantageous to them in a natural context. In the experiment described by Bruce et al., a distractor presented to the ALEs would cause a gaze shift of the AMEs, even when these were already scanning a previous stimulus, but only consistently if the relevance of the new distractor was higher than the primary target. This demonstrates that ALEs maintain significant attentional resources even when AMEs are in use. Given our results and the wider literature, we believe that jumping spiders have employed an ingenious solution to their limited brain resources, by splitting attentive resources between the two eye systems. However, they maintain synergistic coordination, to avoid wasting resources in attentional overlaps. What remains unknown is to what extent these two systems remain separate and how much communication is instead possible between the two.

## Notes

### Competing Interest Statement

The authors have declared no competing interest.

### Summary of Updates

Not all in-text references were reported in the reference list. This version has been corrected.

## References

Blest, A. D., Hardie, R. C., McIntyre, P. and Williams, D. S. (1981). The spectral sensitivities of identified receptors and the function of retinal tiering in the principal eyes of a jumping spider. J. Comp. Physiol. 145, 227–239.

Bolker, B. M., Brooks, M. E., Clark, C. J., Geange, S. W., Poulsen, J. R., Stevens, M. H. H. and White, J.-S. S. (2009). Generalized linear mixed models: a practical guide for ecology and evolution. Trends Ecol. Evol. 24, 127–135.

Bowman, E. M., Brown, V. J., Kertzman, C., Schwarz, U. and Robinson, D. L. (1993). Covert orienting of attention in macaques. I. Effects of behavioral context. J. Neurophysiol. 70, 431–443.

Bradski, G. (2000). The OpenCV Library. Dr Dobbs J. Softw. Tools.

Brooks, M. E., Kristensen, K., Benthem, K. J. van Magnusson, A., Berg, C. W., Nielsen, A., Skaug, H. J., Maechler, M. and Bolker, B. M. (2017). glmmTMB Balances Speed and Flexibility Among Packages for Zero-inflated Generalized Linear Mixed Modeling. R J. 9, 378–400.

Bruce, M., Daye, D., Long, S. M., Winsor, A. M., Menda, G., Hoy, R. R. and Jakob, E. M. (2021). Attention and distraction in the modular visual system of a jumping spider. J. Exp. Biol. 224,.

Bushnell, M. C., Goldberg, M. E. and Robinson, D. L. (1981). Behavioral enhancement of visual responses in monkey cerebral cortex. I. Modulation in posterior parietal cortex related to selective visual attention. J. Neurophysiol. 46, 755–772.

Cross, F. R., Carvell, G. E., Jackson, R. R. and Grace, R. C. (2020). Arthropod Intelligence? The Case for Portia. Front. Psychol. 11,.

De Agrò, M. (2020). SPiDbox: design and validation of an open-source “Skinner-box” system for the study of jumping spiders. J. Neurosci. Methods 346, 108925.

De Agrò, M., Regolin, L. and Moretto, E. (2017). Visual Discrimination Learning in the Jumping Spider Phidippus regius. Anim. Behav. Cogn. 4, 413–424.

De Agrò, M., Rößler, D. C., Kim, K. and Shamble, P. S. (2021). Perception of biological motion by jumping spiders. PLOS Biol. 19, e3001172.

Dolev, Y. and Nelson, X. J. (2014). Innate Pattern Recognition and Categorization in a Jumping Spider. PLoS ONE 9, e97819.

Dolev, Y. and Nelson, X. (2016). Biological relevance affects object recognition in jumping spiders. N. Z. J. Zool. 43, 42–53.

Duelli, P. (1978). Movement detection in the posterolateral eyes of jumping spiders (Evarcha arcuata, Salticidae). J. Comp. Physiol. A 124, 15–26.

Eckstein, M. P., Mack, S. C., Liston, D. B., Bogush, L., Menzel, R. and Krauzlis, R. J. (2013). Rethinking human visual attention: Spatial cueing effects and optimality of decisions by honeybees, monkeys and humans. Vision Res. 85, 5–19.

Evans, K. K., Horowitz, T. S., Howe, P., Pedersini, R., Reijnen, E., Pinto, Y., Kuzmova, Y. and Wolfe, J. M. (2011). Visual attention. WIREs Cogn. Sci. 2, 503–514.

Fenk, L. M. and Schmid, A. (2010). The orientation-dependent visual spatial cut-off frequency in a spider. J. Exp. Biol. 213, 3111–3117.

Foelix, R. F. (2011). Biology of spiders. 3rd ed. Oxford ; New York: Oxford University Press.

Fox, J. and Weisberg, S. (2019). An R Companion to Applied Regression. Third. Thousand Oaks CA: Sage.

Gabay, S., Leibovich, T., Ben-Simon, A., Henik, A. and Segev, R. (2013). Inhibition of return in the archer fish. Nat. Commun. 4, 1657.

Golla, H., Ignashchenkova, A., Haarmeier, T. and Thier, P. (2004). Improvement of visual acuity by spatial cueing: a comparative study in human and non-human primates. Vision Res. 44, 1589–1600.

Greenfield, M. D. and Rand, A. S. (2000). Frogs Have Rules: Selective Attention Algorithms Regulate Chorusing in Physalaemus pustulosus (Leptodactylidae). Ethology 106, 331–347.

Harland, D. P. and Jackson, R. R. (2000). “Eight-legged cats” and how they see - a review of recent research on jumping spiders (Araneae: Salticidae). 11.

Harland, Li, and Jackson (2012). How Jumping Spiders See the World.

Hartig, F. (2018). DHARMa: Residual Diagnostics for Hierarchical (Multi-Level / Mixed) Regression Models.

Humphrey, B., Helton, W. S., Bedoya, C., Dolev, Y. and Nelson, X. J. (2018). Psychophysical investigation of vigilance decrement in jumping spiders: overstimulation or understimulation? Anim. Cogn. 21, 787–794.

Humphrey, B., Helton, W. S. and Nelson, X. J. (2019). Caffeine affects the vigilance decrement of Trite planiceps jumping spiders (Salticidae). J. Comp. Psychol. 133, 551–557.

Hunt, A. R. and Kingstone, A. (2003a). Inhibition of return: Dissociating attentional and oculomotor components. J. Exp. Psychol. Hum. Percept. Perform. 29, 1068–1074.

Hunt, A. R. and Kingstone, A. (2003b). Covert and overt voluntary attention: linked or independent? Cogn. Brain Res. 18, 102–105.

Jackson, R. R. and Cross, F. R. (2011). Spider Cognition. In Advances in Insect Physiology, pp. 115–174. Elsevier.

Jakob, E. M., Long, S. M., Harland, D. P., Jackson, R. R., Carey, A., Searles, M. E., Porter, A. H., Canavesi, C. and Rolland, J. P. (2018). Lateral eyes direct principal eyes as jumping spiders track objects. Curr. Biol. 28, R1092–R1093.

Johnen, A., Wagner, H. and Gaese, B. H. (2001). Spatial Attention Modulates Sound Localization in Barn Owls. J. Neurophysiol. 85, 1009–1012.

Kouider, S. and Dehaene, S. (2007). Levels of processing during non-conscious perception: a critical review of visual masking. Philos. Trans. R. Soc. B Biol. Sci. 362, 857–875.

Land, M. F. (1969a). Movements of the retinae of jumping spiders (Salticidae: dendryphantinae) in response to visual stimuli. J. Exp. Biol. 51, 471–493.

Land, M. F. (1969b). Structure of the Retinae of the Principal Eyes of Jumping Spiders (Salticidae: Dendryphantinae) in Relation to Visual Optics. J. Exp. Biol. 51, 443–470.

Land, M. F. (1985). The Morphology and Optics of Spider Eyes. In Neurobiology of Arachnids (ed. Barth, F. G.), pp. 53–78. Berlin, Heidelberg: Springer Berlin Heidelberg.

Lazareva, O. F., Shimizu, T. and Wasserman, E. A. (2012). How Animals See the WorldComparative Behavior, Biology, and Evolution of Vision. Oxford University Press.

Lenth, R. (2018). emmeans: Estimated Marginal Means, aka Least-Squares Means.

Marote, C. F. O. and Xavier, G. F. (2011). Endogenous-like orienting of visual attention in rats. Anim. Cogn. 14, 535–544.

Melrose, A., Nelson, X. J., Dolev, Y. and Helton, W. S. (2018). Vigilance all the way down: Vigilance decrement in jumping spiders resembles that of humans. Q. J. Exp. Psychol. 2006 747021818798743.

Menda, G., Shamble, P. S., Nitzany, E. I., Golden, J. R. and Hoy, R. R. (2014). Visual Perception in the Brain of a Jumping Spider. Curr. Biol. 24, 2580–2585.

Moore, R. J. D., Taylor, G. J., Paulk, A. C., Pearson, T., van Swinderen, B. and Srinivasan, M. V. (2014). FicTrac: A visual method for tracking spherical motion and generating fictive animal paths. J. Neurosci. Methods 225, 106–119.

Morawetz, L. and Spaethe, J. (2012). Visual attention in a complex search task differs between honeybees and bumblebees. J. Exp. Biol. 215, 2515–2523.

Morehouse, N. (2020). Spider vision. Curr. Biol. 30, R975–R980.

Oliphant, T. E. (2006). A guide to NumPy. Trelgol Publishing USA.

Parker, M. O., Gaviria, J., Haigh, A., Millington, M. E., Brown, V. J., Combe, F. J. and Brennan, C. H. (2012). Discrimination reversal and attentional sets in zebrafish (Danio rerio). Behav. Brain Res. 232, 264–268.

Peirce, J., Gray, J. R., Simpson, S., MacAskill, M., Höchenberger, R., Sogo, H., Kastman, E. and Lindeløv, J. K. (2019). PsychoPy2: Experiments in behavior made easy. Behav. Res. Methods 51, 195–203.

Portugal, R. D. and Svaiter, B. F. (2011). Weber-Fechner Law and the Optimality of the Logarithmic Scale. Minds Mach. 21, 73–81.

Posner, M. I. (2016). Orienting of Attention: Then and Now. Q. J. Exp. Psychol. 69, 1864–1875.

Posner, M. I., Snyder, C. R. and Davidson, B. J. (1980). Attention and the detection of signals. J. Exp. Psychol. Gen. 109, 160–174.

R Core Team (2021). R: A Language and Environment for Statistical Computing. Vienna, Austria: R Foundation for Statistical Computing.

Rizzolatti, G. (1983). Mechanisms of Selective Attention in Mammals. In Advances in Vertebrate Neuroethology (ed. Ewert, J.-P.), Capranica, R. R.), and Ingle, D. J.), pp. 261–297. Boston, MA: Springer US.

Rößler, D. C., De Agrò, M., Kim, K. and Shamble, P. S. (2022a). Static visual predator recognition in jumping spiders. Funct. Ecol. 36, 561–571.

Rößler, D. C., Kim, K., De Agrò, M., Jordan, A., Galizia, C. G. and Shamble, P. S. (2022b). Regularly occurring bouts of retinal movements suggest an REM sleep–like state in jumping spiders. Proc. Natl. Acad. Sci. 119, e2204754119.

Schutten, G.-J., Chan, C., Leeper, T. J., Foster, J. and contributors, and other (2020). readODS: Read and Write ODS Files.

Shimp, C. P. and Friedrich, F. J. (1993). Behavioral and computational models of spatial attention. J. Exp. Psychol. Anim. Behav. Process. 19, 26–37.

Spano, L., Long, S. M. and Jakob, E. M. (2012). Secondary eyes mediate the response to looming objects in jumping spiders (Phidippus audax, Salticidae). Biol. Lett. 8, 949–951.

Sridharan, D., Schwarz, J. S. and Knudsen, E. I. (2014). Selective attention in birds. Curr. Biol. 24, R510–R513.

Steinhoff, P. O. M., Uhl, G., Harzsch, S. and Sombke, A. (2020). Visual pathways in the brain of the jumping spider Marpissa muscosa. J. Comp. Neurol. 528, 1883– 1902.

Strausfeld, N. J. and Barth, F. G. (1993). Two visual systems in one brain: Neuropils serving the secondary eyes of the spider Cupiennius salei. J. Comp. Neurol. 328, 43–62.

Strausfeld, N. J., Weltzien, P. and Barth, F. G. (1993). Two visual systems in one brain: Neuropils serving the principal eyes of the spider Cupiennius salei. J. Comp. Neurol. 328, 63–75.

Tárano, Z. (2015). Choosing a Mate in a Cocktail Party-Like Situation: The Effect of Call Complexity and Call Timing between Two Rival Males on Female Mating Preferences in the Túngara Frog Physalaemus pustulosus. Ethology 121, 749–759.

Theeuwes, J., Kramer, A. F., Hahn, S. and Irwin, D. E. (1998). Our Eyes do Not Always Go Where we Want Them to Go: Capture of the Eyes by New Objects. Psychol. Sci. 9, 379–385.

Ushey, K., Allaire, J. J. and Tang, Y. (2021). reticulate: Interface to “Python.”

van der Walt, S., Colbert, S. C. and Varoquaux, G. (2011). The NumPy Array: A Structure for Efficient Numerical Computation. Comput. Sci. Eng. 13, 22–30.

Van Rossum, G. and Drake, F. L. (2009). Python 3 Reference Manual. Scotts Valley, CA: CreateSpace.

Wagner, U., Baker, L. and Rostron, C. (2014). Searching for inhibition of return in the rat using the covert orienting of attention task. Anim. Cogn. 17, 1121–1135.

Wickham, H. (2009). ggplot2: Elegant Graphics for Data Analysis. Springer-Verlag New York.

Wiederman, S. D. and O’Carroll, D. C. (2013). Selective Attention in an Insect Visual Neuron. Curr. Biol. 23, 156–161.

Winsor, A. M., Pagoti, G. F., Daye, D. J., Cheries, E. W., Cave, K. R. and Jakob, E. M. (2021). What gaze direction can tell us about cognitive processes in invertebrates. Biochem. Biophys. Res. Commun. 564, 43–54.

Yantis, S. and Jonides, J. (1984). Abrupt visual onsets and selective attention: Evidence from visual search. J. Exp. Psychol. Hum. Percept. Perform. 10, 601–621.

Zurek, D. B. and Nelson, X. J. (2012). Saccadic tracking of targets mediated by the anterior-lateral eyes of jumping spiders. J. Comp. Physiol. A 198, 411–417.

Zurek, D. B., Taylor, A. J., Evans, C. S. and Nelson, X. J. (2010). The role of the anterior lateral eyes in the vision-based behaviour of jumping spiders. J. Exp. Biol. 213, 2372–2378.

Zurek, D. B., Cronin, T. W., Taylor, L. A., Byrne, K., Sullivan, M. L. G. and Morehouse, N. I. (2015). Spectral filtering enables trichromatic vision in colorful jumping spiders. Curr. Biol. 25, R403–R404.

