## Supplemental Material 02 for "Separate attentional processes in the two visual systems of jumping spiders": ESM02_full-analysis.html


### ESM02 - Full Analysis

Abstract

This supplement provides the entire R script and output of the
statistical analysis we performed and figures produced, in their
original form. It is presented in the spirit of open and transparent
science, but has not been carefully curated.

### Setup

#### Prepare R environment

```
library(glmmTMB) #for mixed models
library(car) #for anova on mixed models
library(DHARMa) #for goodness of fit of the model
library(emmeans) #for post hoc
library(ggplot2) #to plot
library(reticulate)

use_python('/home/massimodeagro/PycharmProjects/DataAnalysis/venv/bin/python')
```

#### Prepare Python environment

```
import pandas as pd
import os
import matplotlib.pyplot as plt
import numpy as np
import seaborn as sns
```

#### Load data

Will load as separate data frames the two experiments

```
exp1 <- read.csv(paste0(path,'ESM01_Exp1.csv'))
exp2 <- read.csv(paste0(path,'ESM01_Exp2.csv'))


exp1$date <- as.factor(exp1$date)
exp1$subj <- as.factor(exp1$subj)
exp1$sex <- as.factor(exp1$sex)
exp1$trialn <- as.factor(exp1$trialn)
exp1$cond <- as.factor(exp1$cond)
exp1$cuepos <- as.factor(exp1$cuepos)
exp1$targetpos <- as.factor(exp1$targetpos)
exp1$turnside <- as.factor(exp1$turnside)
exp1$congruency <- as.factor(exp1$congruency)

exp2$date <- as.factor(exp2$date)
exp2$subj <- as.factor(exp2$subj)
exp2$sex <- as.factor(exp2$sex)
exp2$trialn <- as.factor(exp2$trialn)
exp2$cond <- as.factor(exp2$cond)
exp2$cuepos <- as.factor(exp2$cuepos)
exp2$targetpos <- as.factor(exp2$targetpos)
exp2$turnside <- as.factor(exp2$turnside)
exp2$congruency <- as.factor(exp2$congruency)
```

```
exp1 = pd.read_csv(path+'ESM01_Exp1.csv')
exp2 = pd.read_csv(path+'ESM01_Exp2.csv')
```

### Analysis

#### Experiment 1 - Spatial cueing task with peripheral cue.

In this experiment, we presented spiders with two focal locations,
30° to the left or to the right of the center of the spiders’ visual
fields. Cues consisted of flashing black and white stimuli, while the
target consisted of a 4° wide circle wobbling up and down. Three
conditions were performed: one in which only the cue appeared, one with
only the target, the last with both. The time passing between target and
cue appearance changed randomly between 0 and 5 seconds, with a peak
frequency of 0.2. the time steps varied logarithmically: a delay of 0.2
had double the chance of appearing in respect to a 0.4 delay.

```
summary(exp1$delay)
```

```
##    Min. 1st Qu.  Median    Mean 3rd Qu.    Max.    NA's 
##  0.0001  0.4124  0.8914  1.1276  1.6191  4.3914    1368
```

```
sns.distplot(exp1['delay'])
```

before proceeding with the main analysis concerning the delay and
congruency effects, we will consider a few general observations

##### Preliminary analyses

###### 1 - Pivot curve description

being this the first usage of this scoring procedure, and
specifically the first time we are converting the delta values produced
by fictrac into binomial data, we need to check how successful our
selection algorithm has been. To do so, we will look at the average
rotational magnitude and duration, and eventually get rid of curves that
result too different to the others or too extreme to be realistic

```
sns.distplot(exp1['peakduration'])
```

```
sns.distplot(exp1['magnitude'])
```

given the selection procedure operated on the raw data, the time span
that defines successful curves has a high consistency, between 0.2 and 1
second, which is realistic to the spider behaviour. The rotation
magnitude instead has some outliers, with almost a 180° rotation. Since
they are a very small number, I will keep them in the data. my guess is
that they actually are two rotations combined, when a spider tries to
pivot towards a stimulus and then immediately repeat the behaviour,
since the visual feedback for the rotation is absent.

###### 2 - response probability across the experiment

Spiders underwent three successive trials, with 12 stimuli per trial.
It is reasonable to assume that across time the response probability may
decrease. This is not crucial for the experiment in question, as these
variables are balanced, but it is still an interesting observation.

```
pmprob <- glmmTMB(turnedtostim~trialn*stimn+ (1|subj), data = exp1, family = binomial())
simres <- simulateResiduals(pmprob)
plot(simres)
```

the fit is good, will proceed with the analysis

```
Anova(pmprob)
```

```
## Analysis of Deviance Table (Type II Wald chisquare tests)
## 
## Response: turnedtostim
##                Chisq Df Pr(>Chisq)    
## trialn       32.9466  2   7.01e-08 ***
## stimn        13.9395  1  0.0001888 ***
## trialn:stimn  0.3499  2  0.8394809    
## ---
## Signif. codes:  0 '***' 0.001 '**' 0.01 '*' 0.05 '.' 0.1 ' ' 1
```

```
e<- emmeans(pmprob, ~trialn)
```

```
## NOTE: Results may be misleading due to involvement in interactions
```

```
pairs(e, adjust='bonferroni')
```

```
##  contrast          estimate    SE  df z.ratio p.value
##  trialn1 - trialn2   0.0139 0.124 Inf   0.112  1.0000
##  trialn1 - trialn3   0.7069 0.138 Inf   5.131  <.0001
##  trialn2 - trialn3   0.6930 0.137 Inf   5.070  <.0001
## 
## Results are given on the log odds ratio (not the response) scale. 
## P value adjustment: bonferroni method for 3 tests
```

```
et <- emtrends(pmprob,~1, var="stimn")
test(et)
```

```
##  1       stimn.trend     SE  df z.ratio p.value
##  overall     -0.0556 0.0153 Inf  -3.633  0.0003
## 
## Results are averaged over the levels of: trialn
```

There is indeed both an effect of trial and one of stimulus number.
will make a plot to observe this effect, for completeness.

```
ggplot(exp1, aes(x=stimn, y=turnedtostim, color=trialn))+
geom_jitter(width = 0.5, height = 0.05)+
geom_smooth(method=glm)
```

```
## `geom_smooth()` using formula = 'y ~ x'
```

###### 3 - sex differences

We now want to know if spiders of different sex have a different
response rate. We expect to find one, because in the literature females
are generally more responsive. We will start by describing our
sample.

```
summary(exp1$sex[!duplicated(exp1$subj)])
```

```
##  f  m 
## 57  9
```

we have many more females over males. the observed difference may be
an effect of this disparity. Regardless, we will proceed to describe our
data

```
pmsex <- glmmTMB(turnedtostim~sex+ (1|subj), data = exp1, family = binomial())
simres <- simulateResiduals(pmsex)
plot(simres)
```

the fit is good, we will proceed with the analysis

```
Anova(pmsex)
```

```
## Analysis of Deviance Table (Type II Wald chisquare tests)
## 
## Response: turnedtostim
##      Chisq Df Pr(>Chisq)  
## sex 2.7238  1    0.09886 .
## ---
## Signif. codes:  0 '***' 0.001 '**' 0.01 '*' 0.05 '.' 0.1 ' ' 1
```

we did found a difference. We proceed with a post-hoc

```
esex <- emmeans(pmsex, ~sex, type = 'response')
pairs(esex)
```

```
##  contrast odds.ratio    SE  df null z.ratio p.value
##  f / m         0.522 0.206 Inf    1  -1.650  0.0989
## 
## Tests are performed on the log odds ratio scale
```

```
esex
```

```
##  sex  prob     SE  df asymp.LCL asymp.UCL
##  f   0.285 0.0331 Inf     0.225     0.354
##  m   0.433 0.0901 Inf     0.271     0.611
## 
## Confidence level used: 0.95 
## Intervals are back-transformed from the logit scale
```

it seems that males respond on average more than females. However,
the limited number of males makes us unable to get to any definitive
conclusion.

###### 4 - directional bias

It is possible that spiders possess a side bias. Again, this would
not pose any significant effect on the experiment, as the positions of
the target are randomized, but it may still be interesting to
observe

```
exp1$turnside_bin[exp1$turnside=='left'] <- 1
exp1$turnside_bin[exp1$turnside=='right'] <- 0

pmside <- glmmTMB(turnside_bin~1+ (1|subj), data = exp1, family = binomial())
simres <- simulateResiduals(pmside)
plot(simres)
```

```
eside <- emmeans(pmside, ~1, type='response')
test(eside)
```

```
##  1       prob     SE  df null z.ratio p.value
##  overall  0.5 0.0185 Inf  0.5   0.000  1.0000
## 
## Tests are performed on the logit scale
```

There is no significant side bias

##### Main Analysis

now to the main analysis. This will be split in two part: first we
will observe the differences between conditions, and then we will
observe the effect of cue on target

###### 1 - Differences between conditions

first, in turn probability. We will maintain stimn and trialn since
we observed them to have an effect

```
mmdiff1 <- glmmTMB(turnedtostim~cond*stimn*trialn+ (trialn|subj), data = exp1, family = binomial(),control=glmmTMBControl(optCtrl = list(iter.max = 30000, eval.max = 40000)))
simres <- simulateResiduals(mmdiff1)
plot(simres)
```

```
Anova(mmdiff1)
```

```
## Analysis of Deviance Table (Type II Wald chisquare tests)
## 
## Response: turnedtostim
##                      Chisq Df Pr(>Chisq)    
## cond              127.7128  2    < 2e-16 ***
## stimn              19.1630  1    1.2e-05 ***
## trialn              8.3800  2    0.01515 *  
## cond:stimn          0.4356  2    0.80428    
## cond:trialn         6.5844  4    0.15955    
## stimn:trialn        0.6309  2    0.72945    
## cond:stimn:trialn   5.0942  4    0.27777    
## ---
## Signif. codes:  0 '***' 0.001 '**' 0.01 '*' 0.05 '.' 0.1 ' ' 1
```

```
ediff <- emmeans(mmdiff1, ~cond*trialn, type='response')
```

```
## NOTE: Results may be misleading due to involvement in interactions
```

```
ediff
```

```
##  cond   trialn   prob      SE  df asymp.LCL asymp.UCL
##  both   1      0.4875 0.08174 Inf   0.33379    0.6437
##  cue    1      0.0156 0.00856 Inf   0.00526    0.0451
##  target 1      0.6313 0.07760 Inf   0.47113    0.7670
##  both   2      0.5162 0.08672 Inf   0.35074    0.6782
##  cue    2      0.0254 0.01303 Inf   0.00922    0.0681
##  target 2      0.4151 0.08554 Inf   0.26245    0.5861
##  both   3      0.2900 0.09474 Inf   0.14219    0.5016
##  cue    3      0.0279 0.01503 Inf   0.00963    0.0784
##  target 3      0.2196 0.07898 Inf   0.10236    0.4098
## 
## Confidence level used: 0.95 
## Intervals are back-transformed from the logit scale
```

```
contrast(ediff, list("Trial1_BothVsCue" = c(1,-1,0,0,0,0,0,0,0),
                     "Trial1_BothVsTarget" = c(1,0,-1,0,0,0,0,0,0),
                     "Trial1_CueVsTarget" = c(0, 1,-1,0,0,0,0,0,0),
                     "Trial2_BothVsCue" = c(0,0,0,1,-1,0,0,0,0),
                     "Trial2_BothVsTarget" = c(0,0,0,1,0,-1,0,0,0),
                     "Trial2_CueVsTarget" = c(0,0,0,0, 1,-1,0,0,0),
                     "Trial3_BothVsCue" = c(0,0,0,0,0,0,1,-1,0),
                     "Trial3_BothVsTarget" = c(0,0,0,0,0,0,1,0,-1),
                     "Trial3_CueVsTarget" = c(0,0,0,0,0,0,0, 1,-1),
                     "All_BothVsCue" = c(0.33,-0.33,0,0.33,-0.33,0,0.33,-0.33,0),
                     "All_BothVsTarget" = c(0.33,0,-0.33,0.33,0,-0.33,0.33,0,-0.33),
                     "All_CueVsTarget" = c(0,0.33,-0.33,0,0.33,-0.33,0,0.33,-0.33)), adjust = "bonferroni")
```

```
##  contrast            odds.ratio       SE  df null z.ratio p.value
##  Trial1_BothVsCue      60.19735 37.64053 Inf    1   6.553  <.0001
##  Trial1_BothVsTarget    0.55559  0.25392 Inf    1  -1.286  1.0000
##  Trial1_CueVsTarget     0.00923  0.00595 Inf    1  -7.270  <.0001
##  Trial2_BothVsCue      40.90345 24.84565 Inf    1   6.110  <.0001
##  Trial2_BothVsTarget    1.50308  0.71083 Inf    1   0.862  1.0000
##  Trial2_CueVsTarget     0.03675  0.02191 Inf    1  -5.541  <.0001
##  Trial3_BothVsCue      14.20786  9.84250 Inf    1   3.831  0.0015
##  Trial3_BothVsTarget    1.45158  0.92183 Inf    1   0.587  1.0000
##  Trial3_CueVsTarget     0.10217  0.07000 Inf    1  -3.330  0.0104
##  All_BothVsCue         31.58462 10.70031 Inf    1  10.191  <.0001
##  All_BothVsTarget       1.06557  0.27369 Inf    1   0.247  1.0000
##  All_CueVsTarget        0.03374  0.01164 Inf    1  -9.823  <.0001
## 
## P value adjustment: bonferroni method for 12 tests 
## Tests are performed on the log odds ratio scale
```

We still observe the effect of stimn, but no interaction. There is an
interaction between trialn and cond, but the post hoc clearly shows that
the pattern remains pretty much the same across trials. The probability
of response decreases as expected, spiders do not turn towards the cue
alone. Indeed, the “cue” stimulus was designed to not give the
impression of being an interesting object, but to just stimulate the
visual field. A flash does not look like a prey or predator, while the
moving dot does, as demonstrated in the literature.

```
mmdiff2 <- glmmTMB(turndelay~cond*stimn*trialn+ (1|trialn*subj), data = exp1, family = gaussian(),
                   control=glmmTMBControl(optCtrl = list(iter.max = 30000, eval.max = 40000)))
simres <- simulateResiduals(mmdiff2)
plot(simres)
```

```
hist(exp1$turndelay, breaks=100)
```

the model fit is not good. Indeed, turndelays are not distributed
normally, but instead they look like a log.

A possible solution, is to model the log of the dependent variable,
rather than the raw value.

```
mmdiff2 <- glmmTMB(log10(turndelay)~cond*stimn*trialn+ (trialn|subj), data = exp1, family = gaussian())
```

```
## Warning in eval(predvars, data, env): Si è prodotto un NaN

## Warning in eval(predvars, data, env): Si è prodotto un NaN
```

```
simres <- simulateResiduals(mmdiff2)
plot(simres)
```

```
hist(log10(exp1$turndelay), breaks=100)
```

```
## Warning in hist(log10(exp1$turndelay), breaks = 100): Si è prodotto un NaN
```

It is better, but still not great. At this point the negative values
are a problem. They exist because in the “both” condition spiders can
turn towards the cue already, but the delay is calculated based on
target. Also, log will regenerate negative numbers for values very near
0. Will produce a cutoff at 0.1

```
exp1$turndelayPos <-exp1$turndelay
exp1$turndelayPos[exp1$turndelayPos<=0.1] <- NA
mmdiff3 <- glmmTMB(log10(turndelayPos)~cond*stimn*trialn+ (trialn|subj), data = exp1, family = gaussian())
simres <- simulateResiduals(mmdiff3)
plot(simres)
```

```
hist(log10(exp1$turndelayPos), breaks=100)
```

Indeed, the model now mostly works.

```
Anova(mmdiff3)
```

```
## Analysis of Deviance Table (Type II Wald chisquare tests)
## 
## Response: log10(turndelayPos)
##                      Chisq Df Pr(>Chisq)    
## cond              377.9517  2  < 2.2e-16 ***
## stimn               0.0503  1   0.822573    
## trialn              0.2821  2   0.868451    
## cond:stimn          5.1782  2   0.075089 .  
## cond:trialn         1.1291  4   0.889631    
## stimn:trialn        7.4397  2   0.024238 *  
## cond:stimn:trialn  14.2110  4   0.006651 ** 
## ---
## Signif. codes:  0 '***' 0.001 '**' 0.01 '*' 0.05 '.' 0.1 ' ' 1
```

```
ediff <- emmeans(mmdiff3, ~cond, type='response')
```

```
## NOTE: Results may be misleading due to involvement in interactions
```

```
ediff
```

```
##  cond   response     SE  df lower.CL upper.CL
##  both      0.562 0.0409 593    0.488    0.649
##  cue       3.730 0.3759 593    3.060    4.546
##  target    0.735 0.0568 593    0.631    0.855
## 
## Results are averaged over the levels of: trialn 
## Confidence level used: 0.95 
## Intervals are back-transformed from the log10 scale
```

```
pairs(ediff)
```

```
##  contrast      ratio     SE  df null t.ratio p.value
##  both / cue    0.151 0.0164 593    1 -17.347  <.0001
##  both / target 0.765 0.0709 593    1  -2.885  0.0113
##  cue / target  5.075 0.5933 593    1  13.897  <.0001
## 
## Results are averaged over the levels of: trialn 
## P value adjustment: tukey method for comparing a family of 3 estimates 
## Tests are performed on the log10 scale
```

Same as before, the delay is significantly higher for the cue
condition over target an both conditions. those last two are also
significantly different. Delays so high in the cue condition suggest
that the observed rotations are just noise: infrequent and strangely
distant from cue apparition. In the both condition, the animals are on
average faster than in the target condition.

These two results together support two hypothesis:

- The cue does not elicit rotations in itself. This makes it safe to
  consider the turns in the “both” condition as being towards the target,
  even if influenced by the cue
- The presence of the cue seem to have an overall effect of decreasing
  detection times. It could however also be that some animals are rotating
  towards the cue, and pull the average down. This however is probably not
  the case as cue alone has such a low probability and long delay.

###### 2 - Effects in the “both” condition

Now we observe the main effect of the experiment. We want to see if
the time passing from appearance of the target to the start of rotation
changes depending on whether the cue is on the same side of the target,
and on the time passing between cue and target appearance.

###### turn delay

the turndelay variable will still be transformed in log. for the same
reason, also the delay passing between cue and target will be
log-transformed

```
allboth <- subset(exp1, exp1$cond == 'both')
allboth <- subset(allboth,allboth$delay>0.1) # For the same reason as before, we don't want to have negative logs too far
allbothfordelay <- subset(allboth, allboth$turnedtostim == 1)


mmdel <- glmmTMB(log10(turndelayPos)~congruency*log10(delay)*trialn*stimn+ (stimn|subj), data = allbothfordelay, family = gaussian())
simres <- simulateResiduals(mmdel)
plot(simres)
```

the model is good enough.

```
Anova(mmdel)
```

```
## Analysis of Deviance Table (Type II Wald chisquare tests)
## 
## Response: log10(turndelayPos)
##                                       Chisq Df Pr(>Chisq)   
## congruency                           7.9905  1   0.004702 **
## log10(delay)                         0.3866  1   0.534100   
## trialn                               0.2974  2   0.861835   
## stimn                                4.9890  1   0.025509 * 
## congruency:log10(delay)              1.8266  1   0.176533   
## congruency:trialn                    0.1551  2   0.925400   
## log10(delay):trialn                  1.3684  2   0.504487   
## congruency:stimn                     0.9634  1   0.326342   
## log10(delay):stimn                   4.5927  1   0.032109 * 
## trialn:stimn                         4.3426  2   0.114027   
## congruency:log10(delay):trialn       0.8793  2   0.644274   
## congruency:log10(delay):stimn        0.1991  1   0.655481   
## congruency:trialn:stimn              0.7823  2   0.676284   
## log10(delay):trialn:stimn            5.3928  2   0.067449 . 
## congruency:log10(delay):trialn:stimn 2.1769  2   0.336741   
## ---
## Signif. codes:  0 '***' 0.001 '**' 0.01 '*' 0.05 '.' 0.1 ' ' 1
```

There is a small effect of congruency. Posthoc follows.

```
edel <- emmeans(mmdel, ~congruency, type='response')
```

```
## NOTE: Results may be misleading due to involvement in interactions
```

```
edel
```

```
##  congruency response     SE  df lower.CL upper.CL
##  0             0.505 0.0277 259    0.454    0.563
##  1             0.584 0.0344 259    0.520    0.656
## 
## Results are averaged over the levels of: trialn 
## Confidence level used: 0.95 
## Intervals are back-transformed from the log10 scale
```

```
pairs(edel)
```

```
##  contrast                  ratio     SE  df null t.ratio p.value
##  congruency0 / congruency1 0.866 0.0432 259    1  -2.894  0.0041
## 
## Results are averaged over the levels of: trialn 
## Tests are performed on the log10 scale
```

the overall delay is smaller in the incongruent condition. will
plot.

```
allboth = exp1[exp1['cond'] == 'both']
allboth = allboth[allboth['delay'] >0.1]
allboth = allboth[allboth['turndelay'] >0.1]
allboth['logturndelay'] = np.log10(allboth['turndelay'])
allboth = allboth[allboth['turnedtostim'] == 1]

sns.violinplot(data=allboth, x='congruency', y='logturndelay')
```

Indeed, very similar. but still there.

So overall, there is no effect of either congruency with delay. This
does not support the hypothesis that jumping spiders are sensitive to
the Posner effect.

Using this methodology however, the Posner effect may not affect the
detection speed, but the detection itself. In human studies, the
subjects are tasked with detecting the target, while spiders do so
spontaneously. The cue may have a rotation probability effect, rather
than a delay

###### turn probability

```
mmdelbin <- glmmTMB(turnedtostim~congruency*log10(delay)*trialn*stimn+ (stimn|subj), data = allboth, family = binomial())

simres <- simulateResiduals(mmdelbin)
plot(simres)
```

```
Anova(mmdelbin)
```

```
## Analysis of Deviance Table (Type II Wald chisquare tests)
## 
## Response: turnedtostim
##                                        Chisq Df Pr(>Chisq)    
## congruency                           17.8217  1  2.426e-05 ***
## log10(delay)                          3.1171  1    0.07747 .  
## trialn                                1.9061  2    0.38556    
## stimn                                 4.3661  1    0.03666 *  
## congruency:log10(delay)               5.2378  1    0.02210 *  
## congruency:trialn                     0.7824  2    0.67624    
## log10(delay):trialn                   2.9417  2    0.22973    
## congruency:stimn                      5.7266  1    0.01671 *  
## log10(delay):stimn                    1.3040  1    0.25349    
## trialn:stimn                          1.2152  2    0.54466    
## congruency:log10(delay):trialn        1.5291  2    0.46554    
## congruency:log10(delay):stimn         1.4052  1    0.23586    
## congruency:trialn:stimn               0.0389  2    0.98073    
## log10(delay):trialn:stimn             0.9140  2    0.63317    
## congruency:log10(delay):trialn:stimn  3.9012  2    0.14219    
## ---
## Signif. codes:  0 '***' 0.001 '**' 0.01 '*' 0.05 '.' 0.1 ' ' 1
```

we found both an effect of congruency and of the delay. there is also
the expected effect of stimn

```
edelbin <- emmeans(mmdelbin, ~ congruency,  type='response')
```

```
## NOTE: Results may be misleading due to involvement in interactions
```

```
edelbin
```

```
##  congruency  prob     SE  df asymp.LCL asymp.UCL
##  0          0.565 0.0718 Inf     0.423     0.697
##  1          0.335 0.0674 Inf     0.218     0.477
## 
## Results are averaged over the levels of: trialn 
## Confidence level used: 0.95 
## Intervals are back-transformed from the logit scale
```

```
pairs(edelbin)
```

```
##  contrast                  odds.ratio    SE  df null z.ratio p.value
##  congruency0 / congruency1       2.58 0.628 Inf    1   3.889  0.0001
## 
## Results are averaged over the levels of: trialn 
## Tests are performed on the log odds ratio scale
```

```
trendstimnbin <- emtrends(mmdelbin, ~ 1, var='stimn', type='response')
```

```
## NOTE: Results may be misleading due to involvement in interactions
```

```
trenddelbin <- emtrends(mmdelbin, ~ congruency, var='delay', type='response')
```

```
## NOTE: Results may be misleading due to involvement in interactions
```

```
trenddelbin
```

```
##  congruency delay.trend    SE  df asymp.LCL asymp.UCL
##  0               -0.108 0.157 Inf    -0.415     0.199
##  1                0.471 0.165 Inf     0.147     0.795
## 
## Results are averaged over the levels of: trialn 
## Confidence level used: 0.95
```

```
test(trenddelbin)
```

```
##  congruency delay.trend    SE  df z.ratio p.value
##  0               -0.108 0.157 Inf  -0.690  0.4903
##  1                0.471 0.165 Inf   2.853  0.0043
## 
## Results are averaged over the levels of: trialn
```

overall, the response probability is much smaller for the congruent
condition. Moreover, while the response is always the same in the
incongruent condition, in the congruent one the response rate is
initially lower, to then go back to normal after 5 second. Will plot

```
allboth = exp1[exp1['cond'] == 'both']
allboth = allboth[allboth['delay'] >0.1]

congruent = allboth[allboth['congruency'] == 1]
congruent['logdelay'] = np.log10(congruent['delay'])
```

```
## <string>:1: SettingWithCopyWarning: 
## A value is trying to be set on a copy of a slice from a DataFrame.
## Try using .loc[row_indexer,col_indexer] = value instead
## 
## See the caveats in the documentation: https://pandas.pydata.org/pandas-docs/stable/user_guide/indexing.html#returning-a-view-versus-a-copy
```

```
incongruent = allboth[allboth['congruency'] == 0]
incongruent['logdelay'] = np.log10(incongruent['delay'])
```

```
## <string>:1: SettingWithCopyWarning: 
## A value is trying to be set on a copy of a slice from a DataFrame.
## Try using .loc[row_indexer,col_indexer] = value instead
## 
## See the caveats in the documentation: https://pandas.pydata.org/pandas-docs/stable/user_guide/indexing.html#returning-a-view-versus-a-copy
```

```
f, ax = plt.subplots(figsize=(7, 7))
sns.regplot(x='logdelay', y='turnedtostim', data=congruent, ax=ax)
sns.regplot(x='logdelay', y='turnedtostim', data=incongruent, ax=ax)
```

#### Experiment 2 - Spatial cueing task with central cue.

This experiment followed a similar procedure to the previous one. We
presented spiders with two focal locations, 30° to the left or to the
right of the center of the spiders’ visual fields. Two types of cues
were presented. The first type consisted of a moving dot, starting from
the center and moving towards either the left or the right focal
location. The second type consisted of a flashing dot, starting from one
side and ending to the other one, occupying 5 distinct location. The
target consisted of a 4° wide circle wobbling up and down. The time
passing between target and cue appearance changed randomly between 0 and
5 seconds, following the same distribution as the first experiment.

before proceeding with the main analysis concerning the delay and
congruency effects, we will consider a few general observations

##### Preliminary analyses

###### 1 - Pivot curve description

For completeness, we will follow the same procedure as before, to
observe and describe the pivots.

```
sns.distplot(exp2['peakduration'])
```

```
sns.distplot(exp2['magnitude'])
```

very similar to experiment 1, which supports the overall validity of
the pivots

###### 2 - response probability across the experiment

As before, we will test for response decrement across stimuli and
trials

```
pmprob <- glmmTMB(turnedtostim~trialn*stimn+ (1|subj), data = exp2, family = binomial())
simres <- simulateResiduals(pmprob)
plot(simres)
```

the fit is good, will proceed with the analysis

```
Anova(pmprob)
```

```
## Analysis of Deviance Table (Type II Wald chisquare tests)
## 
## Response: turnedtostim
##                Chisq Df Pr(>Chisq)    
## trialn        2.6097  1     0.1062    
## stimn        76.4966  1     <2e-16 ***
## trialn:stimn  0.5549  1     0.4563    
## ---
## Signif. codes:  0 '***' 0.001 '**' 0.01 '*' 0.05 '.' 0.1 ' ' 1
```

There is no effect of trial, but an effect of stimulus number. will
make a plot to observe this effect, for completeness.

```
ggplot(exp2, aes(x=stimn, y=turnedtostim, color=trialn))+
geom_jitter(width = 0.5, height = 0.05)+
geom_smooth(method=glm)
```

```
## `geom_smooth()` using formula = 'y ~ x'
```

Indeed, being here only two trials, the difference between them is
lower. in the previous experiment only trial 3 brought the average
down.

###### 3 - sex differences

Moving to sex differences

```
summary(exp2$sex[!duplicated(exp2$subj)])
```

```
##  f  m 
## 58 13
```

we still have many more females over males. the observed difference
may be an effect of this disparity. Regardless, we will proceed to
describe our data

```
pmsex <- glmmTMB(turnedtostim~sex+ (1|subj), data = exp2, family = binomial())
simres <- simulateResiduals(pmsex)
plot(simres)
```

the fit is good, we will proceed with the analysis

```
Anova(pmsex)
```

```
## Analysis of Deviance Table (Type II Wald chisquare tests)
## 
## Response: turnedtostim
##      Chisq Df Pr(>Chisq)  
## sex 3.3935  1    0.06546 .
## ---
## Signif. codes:  0 '***' 0.001 '**' 0.01 '*' 0.05 '.' 0.1 ' ' 1
```

we did not found a difference. We proceed anyway with a post-hoc

```
esex <- emmeans(pmsex, ~sex, type = 'response')
pairs(esex)
```

```
##  contrast odds.ratio    SE  df null z.ratio p.value
##  f / m          2.08 0.826 Inf    1   1.842  0.0655
## 
## Tests are performed on the log odds ratio scale
```

```
esex
```

```
##  sex  prob     SE  df asymp.LCL asymp.UCL
##  f   0.260 0.0355 Inf     0.196     0.335
##  m   0.144 0.0453 Inf     0.076     0.257
## 
## Confidence level used: 0.95 
## Intervals are back-transformed from the logit scale
```

###### 4 - directional bias

About side bias

```
exp2$turnside_bin[exp2$turnside=='left'] <- 1
exp2$turnside_bin[exp2$turnside=='right'] <- 0

pmside <- glmmTMB(turnside_bin~1+ (1|subj), data = exp2, family = binomial())
simres <- simulateResiduals(pmside)
plot(simres)
```

```
eside <- emmeans(pmside, ~1, type='response')
test(eside)
```

```
##  1        prob     SE  df null z.ratio p.value
##  overall 0.535 0.0225 Inf  0.5   1.535  0.1249
## 
## Tests are performed on the logit scale
```

There is no significant side bias

##### Main Analysis

Since both conditions of this experiment contain a cue and a target,
we will mix the two analysis presented in experiment one in a single
model.

###### turn delay

```
exp2onlyturn <- subset(exp2,exp2$delay>0.1) # to remove neg values in log
exp2onlyturn <- subset(exp2onlyturn,exp2onlyturn$turndelay>0.1)
exp2onlyturn <- subset(exp2onlyturn,exp2onlyturn$turnedtostim==1)

mmdel <- glmmTMB(log10(turndelay)~cond*congruency*log10(delay)*stimn+ (trialn|stimn:subj), data = exp2onlyturn, family = gaussian())
simres <- simulateResiduals(mmdel)
plot(simres)
```

```
hist(log10(exp2onlyturn$turndelay), breaks=100)
```

Model’s not perfect, but I doubt it’s getting better than this. will
proceed

```
Anova(mmdel)
```

```
## Analysis of Deviance Table (Type II Wald chisquare tests)
## 
## Response: log10(turndelay)
##                                     Chisq Df Pr(>Chisq)  
## cond                               0.6511  1    0.41973  
## congruency                         4.7559  1    0.02920 *
## log10(delay)                       0.8372  1    0.36021  
## stimn                              0.8799  1    0.34823  
## cond:congruency                    0.2866  1    0.59243  
## cond:log10(delay)                  2.2206  1    0.13618  
## congruency:log10(delay)            0.2268  1    0.63388  
## cond:stimn                         0.2855  1    0.59313  
## congruency:stimn                   1.1690  1    0.27961  
## log10(delay):stimn                 0.6567  1    0.41774  
## cond:congruency:log10(delay)       0.0712  1    0.78965  
## cond:congruency:stimn              1.7733  1    0.18298  
## cond:log10(delay):stimn            0.0092  1    0.92352  
## congruency:log10(delay):stimn      0.0143  1    0.90472  
## cond:congruency:log10(delay):stimn 2.8367  1    0.09213 .
## ---
## Signif. codes:  0 '***' 0.001 '**' 0.01 '*' 0.05 '.' 0.1 ' ' 1
```

There is an effect of congruency.

```
edel <- emmeans(mmdel, ~congruency, type='response')
```

```
## NOTE: Results may be misleading due to involvement in interactions
```

```
edel
```

```
##  congruency response     SE  df lower.CL upper.CL
##  0             0.618 0.0302 372    0.562    0.681
##  1             0.711 0.0373 372    0.641    0.788
## 
## Results are averaged over the levels of: cond 
## Confidence level used: 0.95 
## Intervals are back-transformed from the log10 scale
```

```
pairs(edel, simple='congruency')
```

```
##  contrast                  ratio     SE  df null t.ratio p.value
##  congruency0 / congruency1  0.87 0.0625 372    1  -1.941  0.0530
## 
## Results are averaged over the levels of: cond 
## Tests are performed on the log10 scale
```

```
etdel <- emtrends(mmdel, ~cond, var='log10(delay)', type='response')
```

```
## NOTE: Results may be misleading due to involvement in interactions
```

```
etdel
```

```
##  cond   log10(delay).trend     SE  df lower.CL upper.CL
##  five              -0.0187 0.0679 372  -0.1521    0.115
##  moving             0.0732 0.0498 372  -0.0248    0.171
## 
## Results are averaged over the levels of: congruency 
## Confidence level used: 0.95
```

```
pairs(etdel)
```

```
##  contrast      estimate     SE  df t.ratio p.value
##  five - moving  -0.0919 0.0846 372  -1.087  0.2777
## 
## Results are averaged over the levels of: congruency
```

```
exp2 = exp2[exp2['turnedtostim'] == 1]
exp2 = exp2[exp2['delay'] >=0.1]
exp2['logdelay'] = np.log10(exp2['delay'])

moving = exp2[exp2['cond'] == 'moving']
five = exp2[exp2['cond'] == 'five']

five = five[five['turndelay'] >0.1]
moving = moving[moving['turndelay'] >0.1]
five['logturndelay'] = np.log10(five['turndelay'])
moving['logturndelay'] = np.log10(moving['turndelay'])


f, axs = plt.subplots(1,2,figsize=(7, 7))
sns.violinplot(data=moving,ax=axs[0], x='congruency', y='logturndelay')
sns.violinplot(data=five,ax=axs[1], x='congruency', y='logturndelay')
```

Here the effect, even though still tiny, it’s better appreciable. In
both conditions the spiders are faster in incongruent trials.

We will proceed with the binomial analysis as for experiment 1

###### turn probability

```
mmdelbin <- glmmTMB(turnedtostim~cond*congruency*log10(delay)*stimn+ (trialn:stimn|subj), data = exp2, family = binomial())

simres <- simulateResiduals(mmdelbin)
plot(simres)
```

```
Anova(mmdelbin)
```

```
## Analysis of Deviance Table (Type II Wald chisquare tests)
## 
## Response: turnedtostim
##                                      Chisq Df Pr(>Chisq)    
## cond                               15.6177  1  7.753e-05 ***
## congruency                          0.6137  1    0.43341    
## log10(delay)                        3.5913  1    0.05808 .  
## stimn                              44.4343  1  2.630e-11 ***
## cond:congruency                     0.8976  1    0.34342    
## cond:log10(delay)                   0.6651  1    0.41475    
## congruency:log10(delay)             0.6567  1    0.41772    
## cond:stimn                          4.7173  1    0.02986 *  
## congruency:stimn                    6.1487  1    0.01315 *  
## log10(delay):stimn                  1.7622  1    0.18435    
## cond:congruency:log10(delay)        0.0111  1    0.91625    
## cond:congruency:stimn               0.2509  1    0.61648    
## cond:log10(delay):stimn             0.6836  1    0.40836    
## congruency:log10(delay):stimn       0.1201  1    0.72893    
## cond:congruency:log10(delay):stimn  0.1878  1    0.66473    
## ---
## Signif. codes:  0 '***' 0.001 '**' 0.01 '*' 0.05 '.' 0.1 ' ' 1
```

There is a difference between conditions, and the expected effect of
stimn. There is a curious interaction between condition and stimn, and
congruency and stimn that I will try to further explore. Let’s start
with the main cond effect.

```
edelbin <- emmeans(mmdelbin, ~ cond,  type='response')
```

```
## NOTE: Results may be misleading due to involvement in interactions
```

```
edelbin
```

```
##  cond    prob     SE  df asymp.LCL asymp.UCL
##  five   0.086 0.0240 Inf    0.0491     0.146
##  moving 0.263 0.0501 Inf    0.1769     0.372
## 
## Results are averaged over the levels of: congruency 
## Confidence level used: 0.95 
## Intervals are back-transformed from the logit scale
```

```
pairs(edelbin)
```

```
##  contrast      odds.ratio     SE  df null z.ratio p.value
##  five / moving      0.264 0.0784 Inf    1  -4.482  <.0001
## 
## Results are averaged over the levels of: congruency 
## Tests are performed on the log odds ratio scale
```

Indeed, the response rate for the moving dot condition is double the
one for the five.

```
trenddelbin <- emtrends(mmdelbin, ~ cond*congruency, var='stimn', type='response')
```

```
## NOTE: Results may be misleading due to involvement in interactions
```

```
trenddelbin
```

```
##  cond   congruency stimn.trend     SE  df asymp.LCL asymp.UCL
##  five   0               -0.281 0.0649 Inf    -0.408   -0.1536
##  moving 0               -0.156 0.0547 Inf    -0.263   -0.0491
##  five   1               -0.436 0.0707 Inf    -0.574   -0.2974
##  moving 1               -0.250 0.0560 Inf    -0.360   -0.1407
## 
## Confidence level used: 0.95
```

```
contrast(trenddelbin, list(FiveVsMoving = c(0.5,-0.5,0.5,-0.5),
                           IncongruentVsCongruent = c(0.5,0.5,-0.5,-0.5)), adjust='bonferroni')
```

```
##  contrast               estimate     SE  df z.ratio p.value
##  FiveVsMoving             -0.155 0.0663 Inf  -2.340  0.0386
##  IncongruentVsCongruent    0.125 0.0496 Inf   2.514  0.0239
## 
## P value adjustment: bonferroni method for 2 tests
```

It seems that across stimuli, the response rate drops much faster in
the five condition over the moving condition, and in the congruent
condition over the incongruent one.
