## Supplemental Material 02 for "Separate attentional processes in the two visual systems of jumping spiders": ESM02_scoringExp1.py

```
"""
Experiment 1 - Attentive priming with peripheral cues
This script takes FicTrac output and transform it into the dataset analyzed in the paper.
This is done by: (the steps are numbered across the script as well)

    1. opening one single-trial .dat from fictrac and the .csv with timestamps.
    2. applying Butterworth filter to all three axis (referenced to the spider point of view)
    3. Finding z-only (or almost z-only) movements, by subtracting absolute value of x and y to z,
       This way, we get a z line, the highest the value, the more the movement was purely on the Z axis.
       Z peaks that were lower than X or Y in the same time point get deleted.
    4. save the timed data onto a file as a checkpoint

    ---
    from here, the analysis changes from what described in De Agrò et al 2021. There, we took every
    peak and averaged them out to find preference. Here, we need to infer detection and delay. This means
    we need the one and only rotation (specifically the first). Here's the procedure:
    ---

    5. Load back the timed fictrac and the csv containing the stimuli appearance, position and timing
    6. Cut the data into sections, going from cue appearance to 10 seconds after the disappearance of target
    7. Get z data cumulatively
    8. Compute the second derivative and find peaks, this reveals change points in the data
    9. For every data block, compute regression line.
   10. From every line, select the ones with a minimum inclination (rotation speed) and with a minimum
       difference from start and end (total rotation made) and a maximum span (rotation duration)
   11. Take the first rotation made for every stimulus,
   12. Save. new row into results table, if no rotation has been found, save NA for data.
"""

#### ---- LOAD PACKAGES --- ####

import pandas as pd
import os
import numpy as np
from scipy import signal, stats


#### ---- SET PATH OF FILES --- ####

path = '' #path of the main folder
rawpath = path+'Data_todo/'
timedpath = path+'Data_done/'
resultspath = path+'Results_incremental/'


#### ---- READ ALL DAT FILES IN THE FOLDER ---- ####
files = os.listdir(rawpath)
donefiles = os.listdir(timedpath)
filenames = []
filedatnames =[]
for file in files:
    if file.endswith('.dat'):
        filenames.append(file.split('.')[0][0:-16])
        filedatnames.append(file)


for filename, filedatname in zip(filenames,filedatnames): # For all files
    if filename+'_timedfictrac.csv' not in donefiles:

        ### 1. opening one single-trial .dat from fictrac and the .csv with timestamps.

        time = pd.read_csv(rawpath+filename+'_timestamps.csv',header=None, names=['videotime'])

        fictrac = pd.read_csv(rawpath+filedatname, header=None)
        fictrac.columns = ['framec', 'DELTArvCamX', 'DELTArvCamY', 'DELTArvCamZ',
                     'DELTArvERR',
                     'DELTArvLabX', 'DELTArvLabY', 'DELTArvLabZ',
                     'ABSOLUTErvCamX', 'ABSOLUTErvCamY', 'ABSOLUTErvCamZ',
                     'ABSOLUTErvLabX', 'ABSOLUTErvLabY', 'ABSOLUTErvLabZ',
                     'INTEGRATEDposX', 'INTEGRATEDposY', 'INTEGRATEDheading',
                     'DIR', 'SPEED',
                     'INTEGRATEDmotionX', 'INTEGRATEDmotionY',
                     'timestamp_fictrac', 'seqCounter', 'deltaTime', 'altTime']

        ### 2. applying Butterworth filter to all three axis (referenced to the spider point of view)

        b, a = signal.butter(3, 0.05)
        fictrac['DELTArvLabXfilt'] = signal.filtfilt(b, a, fictrac['DELTArvLabX'].values)
        fictrac['DELTArvLabYfilt'] = signal.filtfilt(b, a, fictrac['DELTArvLabY'].values)
        fictrac['DELTArvLabZfilt'] = signal.filtfilt(b, a, fictrac['DELTArvLabZ'].values)
        fictrac['time'] = fictrac['framec']
        for rown in range(len(time)):
            fictrac.loc[fictrac['framec'] == rown, 'time'] = time['videotime'][rown]
            print(rown)

        ### 3. Finding z-only (or almost z-only) movements, by subtracting absolute value of x and y to z,
        ### This way, we get a z line, the highest the value, the more the movement was purely on the Z axis.
        ### Z values that were lower than X or Y in the same time point get deleted.

        Zclean = []
        for n in range(len(fictrac)):
            z = fictrac['DELTArvLabZfilt'].values[n]
            x = fictrac['DELTArvLabXfilt'].values[n]
            y = fictrac['DELTArvLabYfilt'].values[n]
            #print('z: '+str(z)+' - x: '+str(x)+' - y: '+str(y))
            o = abs(z) - abs(x) - abs(y)

            if o < 0 :
                o = 0
            if z < 0:
                o = o*-1
            Zclean.append(o)
        fictrac['DELTArvLabZclean'] = Zclean
        fictrac['DELTArvLabZAbsolute'] = abs(fictrac['DELTArvLabZclean'])

        ### 4. save the timed data onto a file as a checkpoint

        fictrac.to_csv(timedpath+filename+'_timedfictrac.csv', index=False)


files = os.listdir(timedpath)

filenames = []
filedatnames =[]
for file in files:
    if file.endswith('_timedfictrac.csv'):
        filenames.append(file.split('.')[0][0:-13])
        filedatnames.append(file)

for filename, filedatname in zip(filenames,filedatnames):
    try:

        ### 5. Load back the timed fictrac and the csv containing the stimuli appearance, position and timing

        print(filename)

        # this will contain results
        single_trial_result = [['date','subj','sex','trialn','stimn','cond','cuepos','targetpos','delay', 'turn','turnside','turndelay','magnitude','peakduration']]
        stim = pd.read_csv(rawpath+filename+'_stim.csv')
        date = stim['date'][0]
        subj = stim['name'][0]
        sex = stim['sex'][0]
        trialn = stim['trialn'][0]
        cond = stim['condition'][0]
        if cond == 'b':
            cond = 'both'
        elif cond == 'c':
            cond = 'cue'
        else:
            cond = 'target'

        fictrac = pd.read_csv(timedpath+filedatname)


        stimspeaks = []
        toconsidertime = []
        toconsiderval = []
        peakdelays = []
        cuedelays = []
        for stimn in range(12):
            currstimpeaks = []

            ### 6. Cut the data into sections, going from cue appearance to 10 seconds after the disappearance of target

            if cond == 'both' or cond == 'cue': # for both and cue, start at cue
                stimstart = stim['timestamp'][stim['stimn']== stimn][stim['event']=='start'][stim['stim']=='cue'].item()
                cuestart = stim['timestamp'][stim['stimn'] == stimn][stim['event'] == 'start'][stim['stim'] == 'cue'].item()
            else: #for only target, start at target
                stimstart = stim['timestamp'][stim['stimn'] == stimn][stim['event'] == 'start'][stim['stim'] == 'target'].item()

            if cond == 'both' or cond == 'target': #for both and target, end after 5 seconds of targed disappearance
                stimend = stim['timestamp'][stim['stimn']== stimn][stim['event']=='end'][stim['stim']=='target'].item()
                targetstart = stim['timestamp'][stim['stimn'] == stimn][stim['event'] == 'start'][
                    stim['stim'] == 'target'].item()

            else: #for cue, end after 5 seconds of cue disappearance
                stimend = stim['timestamp'][stim['stimn']== stimn][stim['event']=='end'][stim['stim']=='cue'].item()
                targetstart = stim['timestamp'][stim['stimn'] == stimn][stim['event'] == 'start'][
                    stim['stim'] == 'cue'].item()

            if cond == 'both': # for both, calculate delay between cue and target
                delay = stim['timestamp'][stim['stimn']== stimn][stim['event']=='start'][stim['stim']=='target'].item()-\
                       stim['timestamp'][stim['stimn']== stimn][stim['event']=='end'][stim['stim']=='cue'].item()
            else: #otherwise, no delay
                delay = 'NA'
            if cond == 'cue': #find cue position
                cuepos = stim['pos'][stim['stimn']== stimn][stim['event']=='start'][stim['stim']=='cue'].item()
                targetpos = 'NA'
            elif cond == 'target':  #find target positio
                targetpos = stim['pos'][stim['stimn']== stimn][stim['event']=='start'][stim['stim']=='target'].item()
                cuepos = 'NA'
            else:
                targetpos = stim['pos'][stim['stimn'] == stimn][stim['event'] == 'start'][stim['stim'] == 'target'].item()
                cuepos = stim['pos'][stim['stimn']== stimn][stim['event']=='start'][stim['stim']=='cue'].item()

            # here cut based on start and end
            focustab = fictrac[fictrac['time'] > stimstart]
            focustab = focustab[focustab['time'] < stimend + 10]

            ### 7. Get z data cumulatively

            xcumulative = []
            for val in focustab.DELTArvLabX.values :
                if len(xcumulative) == 0 :
                    xcumulative.append(val)
                else :
                    xcumulative.append(xcumulative[-1] + val)
            ycumulative = []
            for val in focustab.DELTArvLabY.values :
                if len(ycumulative) == 0 :
                    ycumulative.append(val)
                else :
                    ycumulative.append(ycumulative[-1] + val)
            zcumulative = []
            for val in focustab.DELTArvLabZ.values :
                if len(zcumulative) == 0 :
                    zcumulative.append(val)
                else :
                    zcumulative.append(zcumulative[-1] + val)


            ### 8. Compute the second derivative and find peaks, this reveals change points in the data
            if len(zcumulative) > 0 :
                window = 81

                filter = signal.savgol_filter(zcumulative, window_length=window, polyorder=2, deriv=2)
                zfilter = stats.zscore(filter)
                posp, valp = signal.find_peaks(zfilter, prominence=0.1)
                timep = [focustab['time'].values[p] for p in posp]
                posn, valn = signal.find_peaks(zfilter * -1, prominence=0.1)
                timen = [focustab['time'].values[p] for p in posn]

                zbreaks = np.sort(np.concatenate(([0], posp, posn, [len(zcumulative) - 1])))
                ztimebreaks = np.sort(np.concatenate(([stimstart], timep, timen, [stimend + 5])))

                xcumulative = np.rad2deg(xcumulative)
                ycumulative = np.rad2deg(ycumulative)
                zcumulative = np.rad2deg(zcumulative)

                turnsections = []

                ### 9. For every data block, compute regression line.

                for start, end in zip(zbreaks[0 :-1], zbreaks[1 :]) :
                    sectionstarttime = focustab['time'].values[start]
                    sectionendtime = focustab['time'].values[end]

                    # here is regression, will extract values from here below
                    vals = stats.linregress(list(range(start, end)), zcumulative[start :end])
                    slope = vals[0]
                    intc = vals[1]
                    st = intc + slope * start
                    nd = intc + slope * end
                    diff = st - nd
                    if diff > 0 :
                        turnside = 'right'
                    elif diff < 0 :
                        turnside = 'left'
                    else :
                        turnside = None

                    magnitude = abs(diff)
                    timespan = sectionendtime - sectionstarttime
                    deltax = max(xcumulative) - min(xcumulative)
                    deltay = max(ycumulative) - min(ycumulative)

                    # save the regression lines into a dictionary
                    turnsections.append({'index' : {'start' : start, 'end' : end},
                                         'time' : {'start' : sectionstarttime, 'end' : sectionendtime,
                                                   'span' : timespan},
                                         'reg' : {'start' : st, 'end' : nd, 'slope' : slope, 'intercept' : intc,
                                                  'diff' : diff,
                                                  'magnitude' : magnitude},
                                         'turnside' : turnside, 'overlap' : {'x' : deltax, 'y' : deltay}})


                    ### 10. From every line, select the ones with a minimum inclination (rotation speed) and with a minimum
                    ###     difference from start and end (total rotation made) and a maximum span (rotation duration)
                    confirmedturnsections = []
                    lastadded = False
                    for en, section in enumerate(turnsections) :
                        # these nested if are the ever more restrictive conditions to define a rotation
                        if section['time']['span'] > 0.1 and section['time']['span'] < 1.5 :
                            if section['overlap']['x'] < section['reg']['magnitude'] * 0.5 and section['overlap']['y'] < \
                                    section['reg']['magnitude'] * 0.5 :
                                if section['reg']['magnitude'] > 30 / 4 :
                                    if lastadded :
                                        if section['turnside'] != turnsections[en - 1]['turnside'] :
                                            if section['reg']['magnitude'] > turnsections[en - 1]['reg'][
                                                'magnitude'] / 2 :
                                                confirmedturnsections.append(section)
                                                lastadded = True
                                            else :
                                                lastadded = False
                                    else :
                                        confirmedturnsections.append(section)
                                        lastadded = True
                                else :
                                    lastadded = False
                            else :
                                lastadded = False
                        else :
                            lastadded = False


                    ### 11. Take the first rotation made for every stimulus, with direction congruent with the position of the stimulus
                    if len(confirmedturnsections) > 0 :
                        turn = 1
                        turndelay = confirmedturnsections[0]['time']['start'] - targetstart
                        turnside = confirmedturnsections[0]['turnside']

                        peakduration = confirmedturnsections[0]['time']['end'] - confirmedturnsections[0]['time'][
                            'start']
                        peakmagnitude = confirmedturnsections[0]['reg']['magnitude']
                    else :
                        turn = 0
                        turndelay = 'NA'
                        turnside = 'NA'
                        peakduration = 'NA'
                        peakmagnitude = 'NA'
            else:
                turn = 'NA'
                turndelay = 'NA'
                turnside = 'NA'
                peakduration = 'NA'
                peakmagnitude = 'NA'

            ### 12. Save. new row into results table, if no rotation has been found, save NA for data.
            single_trial_result.append([date,subj,sex,trialn,stimn,cond,cuepos,targetpos,delay, turn,turnside,turndelay,peakmagnitude,peakduration])

        # Save single trial file
        np.savetxt(resultspath+filename+'_result.csv',single_trial_result, delimiter=",", fmt='%s')
    except FileNotFoundError:
        print('missing')


### THIS SECTION MERGES ALL FILES
import os

files = os.listdir(resultspath)

tables = []

for n, file in enumerate(files) :
    if file.endswith('result.csv') :
        tables.append(pd.read_csv(resultspath + file))

result = pd.concat(tables)
result.to_csv(resultspath + 'all.csv', index=False)
```
